# Bacteria elicit a phage tolerance response subsequent to infection of their neighbors

**DOI:** 10.1101/2021.02.16.428622

**Authors:** Elhanan Tzipilevich, Osher Pollak-Fiyaksel, Sigal Ben-Yehuda

**Author notes:** To whom correspondence should be addressed. (S. B-Y). These authors contributed equally to this work (Elhanan Tzipilevich, Osher Pollak-Fiyaksel).

## Abstract

Plaque occurrence on a bacterial lawn manifests successive rounds of bacteriophage infection. Yet, mechanisms evolved by bacteria to limit plaque spread have been hardly explored. Here we investigated the dynamics of plaque development by lytic phages infecting the bacterium *Bacillus subtilis*. We report that plaque expansion is followed by a constriction phase owing to bacterial growth into the plaque zone. This phenomenon is caused by an adaptive process, herein termed “phage tolerance response”, elicited by non-infected bacteria located at the plaque rim upon sensing infection of their neighbors. The temporary phage-tolerance is executed by the stress response RNA polymerase sigma factor σ^X^, primarily through activation of the *dlt* operon, encoding enzymes that catalyze D-alanylation of cell wall teichoic acid polymers, the major attachment sites for phages infecting Gram-positive bacteria. D-alanylation impedes phage binding and hence infection, thus enabling the uninfected bacteria to form a protective shield opposing plaque spread.

## Introduction

Plaques, reporting bacterial clearance, are the hallmark of bacteriophage (phage) infection since the pioneering discoveries made by Twort and d’Herelle at the beginning of the 20^th^ century (Chanishvili, 2012; d’Hérelles, 1917; Twort, 1914). Plaques are basically visible holes, formed on a lawn of bacteria grown on a solid surface, that report bacterial clearance following successive cycles of infection, including phage adsorption, replication and spread to nearby hosts. Intriguingly, plaques exhibit a considerable variation in shape according to the host and the infecting phage, and are predominantly restricted in size (Abedon and Yin, 2009). It has been proposed that such size limitation is achieved, at least in part, by the entry of bacteria into stationary phase, which frequently restrains phage replication (Abedon and Yin, 2009). However, although plaque employment as a method for monitoring phage infection began decades ago, relatively little is known about the kinetics of plaque development. Furthermore, factors limiting plaque size and expansion are mostly unrevealed.

In this study, we investigated the dynamics of plaque formation, utilizing the Gram-positive soil bacterium *Bacillus subtilis* (*B. subtilis*) and its lytic phages SPP1 and Phi29. Binding of phages to *B. subtilis* is commonly mediated by wall teichoic acid (WTA) polymers, a diverse family of cell surface glycopolymers containing phosphodiester-linked glycerol repeat units poly(Gro-P), decorated by glucose and D-alanine moieties, and anchored to peptidoglycan (PG) through an N-acetylmannosaminyl (Brown et al., 2013). WTA polymers were found to be crucial surface components, required for invasion by manifold phages into Gram-positive bacteria [e.g. *Bacilli*, *Staphylococci*, *Listeria* (Habusha et al., 2019; Ingmer et al., 2019; Lindberg, 1973; Sumrall et al., 2020)]. SPP1, a double-stranded DNA (dsDNA) phage (44 kb) and a member of the Siphoviridae family, characterized by a long noncontractile tail (Alonso et al., 1997), initiates infection by reversible binding of the tail tip to poly-glycosylated WTA (gWTA). Subsequently, SPP1 binds irreversibly to its membrane receptor protein YueB, resulting in DNA injection into the bacterium cytoplasm (Baptista et al., 2008; Sao-Jose et al., 2004). Phi29 phage, which is significantly smaller (19.3 kb) and belongs to the Podoviridae family, harboring a short noncontractile tail (Salas, 2012), was shown to entirely rely on intact gWTA for infection (Young, 1967). The fact that phages assigned to distinct families utilize gWTA to invade the host strengthens the vital role of these polymers in host recognition by phages and implicates them as major elements for host cell vulnerability. Consistent with this notion, we have shown that a mutant bacteriophage capable of bypassing the need for binding the glucosyl residues, decorating the WTA polymers, gained a broader host range, as it could infect non-host bacterial species presenting dissimilar glycosylation patterns (Habusha et al., 2019).

Previously, we demonstrated that phages could occasionally invade resistant cells that acquire phage receptors from their sensitive neighbors, highlighting the importance of understanding infection dynamics in a temporal and spatial fashion (Tzipilevich et al., 2017). Here, we visualized plaques formed on a lawn of *B. subtilis* bacteria. We revealed that plaque spreading is followed by a phase of plaque constriction mediated by bacterial regrowth into the plaque zone, thereby decreasing the plaque size. Characterization of the plaque constriction phase exposed a temporary immunity mechanism, enabling bacteria to tolerate infection by remodeling WTA polymers. This modification reduces phage binding and restricts phage spread. Unlike other mechanisms affording long-term bacterial immunity to phages, such as restriction enzymes and CRISPR (Labrie et al., 2010), this tolerance mechanism confers a transient adaptive response, providing protection to the uninfected bacterial population subsequent to infection of their neighbors.

## Results

### Plaque expansion is followed by a phase of plaque constriction

To explore the bacterial population dynamic during phage attack on solid surfaces, we followed the process of plaque formation by SPP1 on a lawn of mCherry-labeled *B. subtilis* cells at high resolution, using time lapse confocal microscopy. Maximal SPP1 plaque size was detected approximately eight hours post infection, but remarkably the spread was counteracted by bacteria growing into the plaque area, limiting plaque expansion (Figure 1A; Movie S1). Consequently, the final plaque diameter measured after overnight incubation was significantly smaller than the maximal size reached during plaque development (Figure 1A). To further explore this phenomenon, we followed the kinetics of plaque formation on agar plates, the typical methodology used over the years for estimation of plaque forming units (PFU) [e.g. (Abedon and Yin, 2009; Ellis and Delbruck, 1939)]. Consistent with the confocal microscopy results, plate monitoring revealed a steep expansion phase that was proceeded by a gradual decrease in plaque size, with the final zone being approximately 50% of the maximal plaque area measured during the process (Figure 1B-1C; Movie S2). This plaque constriction occurrence was also evident when bacteria were infected with the distinct lytic phage Phi29 (Figure 1D), indicating that such a kinetic pattern is widespread.

**Figure 1.**
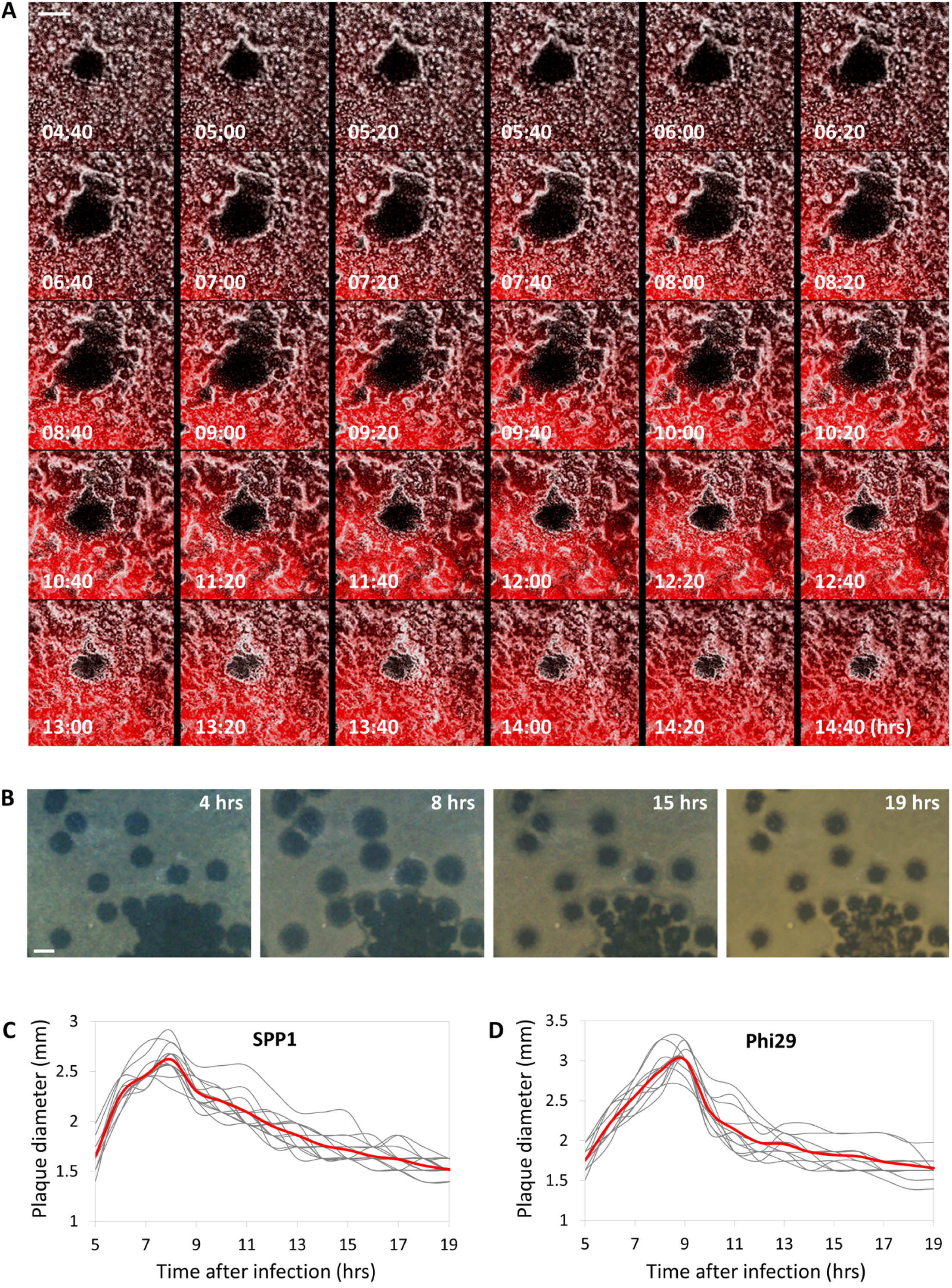
Plaque formation dynamics reveals phases of expansion and constriction. **(A)** BDR2637 (P*_veg_-mCherry*) cells were infected with low concentrations (10^-8^ PFU/ml) of SPP1, placed on an agarose pad and plaque formation was followed by time lapse confocal microscopy. Shown are overlay images from mCherry signal (red) and phase contrast (grey) of the bacterial lawn captured at the indicated time points (hrs). The plaque is seen as a hole formed on the bacterial lawn. Scale bar 150 µm. Corresponds to Movie S1. **(B)** Plaque formation was monitored by automated scanning (Levin-Reisman et al., 2010) on a lawn of PY79 (WT) cells grown on an MB agar plate. Shown are images captured at the indicated time points (hrs). Scale bar 2 mm. Corresponds to Movie S2. **(C-D)** The dynamic of SPP1 (C) and Phi29 (D) plaque formation following PY79 (WT) infection was monitored as described in (B). Shown is the diameter of individual plaques for each phage (n ≥ 12), with the average highlighted in red.

To concomitantly track plaque development and phage localization, we followed plaque formation by SPP1 harboring its lysin gene (*gp51*) fused to a yellow fluorescent protein (YFP) as a sole copy (SPP1-*_lysin-yfp_*) (Tzipilevich et al., 2017). The plaque periphery was occupied by high concentrations of phages, even throughout the phase of plaque constriction (Figure S1; Movie S3), signifying that bacteria at the rim could withstand the presence of phages. Isolating bacteria from the edge of 30 different plaques subsequent to the constriction phase and re-plating them over plate-containing phages revealed the bacteria to remain phage sensitive. We refer to the phenomenon of phage sensitive bacteria that can confront phages at the plaque circumference as “phage tolerance”.

### SigX is necessary for plaque constriction

The phenomenon of plaque constriction directed by phage sensitive bacteria prompted us to postulate that bacteria residing at the plaque periphery could sense a danger signal, emanating from nearby infected bacteria, and in turn mount a transient phage tolerance response. We further reasoned that such a response could be orchestrated by one of the *B. subtilis* extra-cytoplasmic function (ECF) RNA polymerase sigma (σ) factors that are activated in response to stress imposed on the cell envelope (Helmann, 2016). To examine if any of the known sigma factors are required for plaque constriction, we deleted each of the seven ECF sigma factors of *B. subtilis* and assayed their impact on the final plaque size. Intriguingly, *ΔsigX* strain exhibited significantly larger plaques in comparison to wild type (WT) when challenged with SPP1 or Phi29 phages (Figure 2A; Figure S2A). Furthermore, monitoring plaque dynamics revealed that this size difference is due to the substantial attenuation of the plaque constriction phase (Figure 2B-2C). Importantly, *ΔsigX* cells propagated with kinetics similar to that of WT cells (Figure S2B), indicating that the observed phenotype was not due to growth perturbation. To further elucidate the role of SigX in counteracting phage spread, we challenged *ΔsigX* cells with SPP1 or Phi29 phages in liquid cultures. No difference in lysis kinetics was monitored when bacteria were infected at 1:1 (phage:bacteria) multiplicity of infection (MOI). However, while infected with low MOI (phage:bacteria 1:20), *ΔsigX* cells lysed significantly faster than WT cells (Figure 2D; Figure S2C). These results are consistent with the view that uninfected bacteria induce a SigX-regulated defense response, capable of tempering future phage infections.

**Figure 2.**
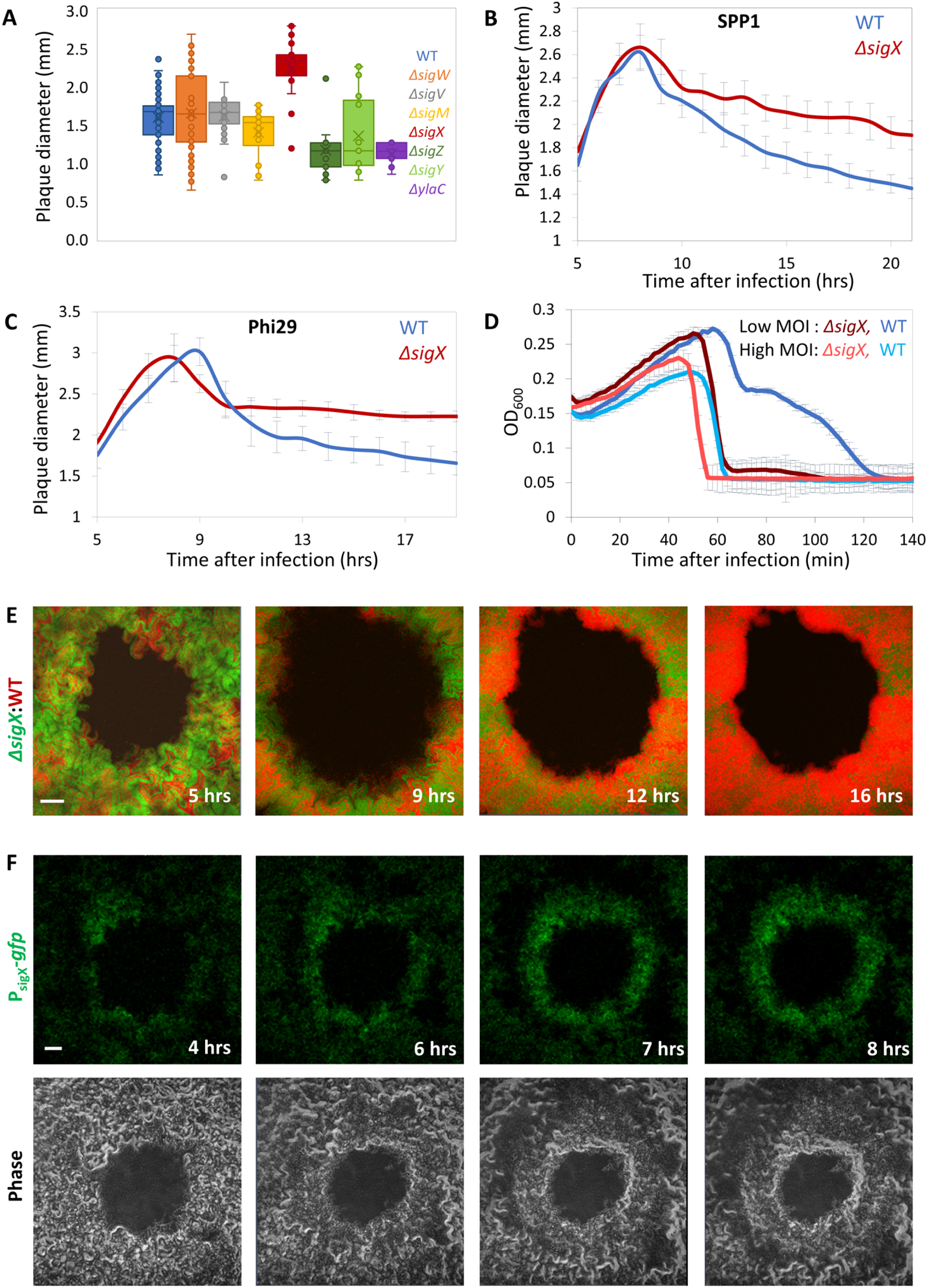
SigX is required for plaque constriction. **(A)** The indicated strains were infected with SPP1 and plaque diameter was monitored after 20 hrs of incubation. Shown is plaque diameter distribution for each strain (n ≥ 54). **(B-C)** Plaque formation dynamic of SPP1 (B) and Phi29 (C) was monitored by automated scanning (Levin-Reisman et al., 2010) on a lawn of PY79 (WT) or ET19 (*ΔsigX*) cells grown on an MB agar plate. Shown are average values and SD from kinetic of random plaques for each strain (n ≥ 7). **(D)** PY79 (WT) and ET19 (*ΔsigX*) cells were infected with SPP1 at either high (phages:bacteria 1:1) or low (1:20) MOI, and OD_600_ was followed at 2 min intervals. Shown is a representative experiment out of 6 biological repeats, and the average values and SD of 8 technical repeats. **(E)** BDR2637 (P_veg_*-mCherry*) (WT, red) and ET191 (P_rrnE_*-gfp*, *ΔsigX*) (*ΔsigX*, green) cells were mixed, infected with low concentrations (10^-8^ PFU/ml) of SPP1, placed on an agarose pad and plaque formation was followed by time lapse confocal microscopy. Shown are overlay images from mCherry (red) and GFP (green) signal of the bacterial lawn captured at the indicated time points (hrs). The plaque is seen as a hole formed on the bacterial lawn. Scale bar 100 µm. Corresponds to Movie S4. **(F)** ET27 (P_sigX_*-gfp*) cells were infected with low concentrations (10^-8^ PFU/ml) of SPP1, placed on an agarose pad, and plaque formation was followed by time lapse confocal microscopy. Shown are fluorescence from GFP (upper panels) and corresponding phase contrast images (lower panels), captured at the indicated time points (hrs). The plaque is seen as a hole formed on the bacterial lawn. Scale bar 100 µm.

To compare the response to phage attack of WT and *ΔsigX* cells in real time, we mixed mCherry-labeled WT cells with GFP-labeled *ΔsigX* cells and monitored plaque dynamics following infection with SPP1, utilizing time lapse confocal microscopy. At the initial stages of phage spreading, both strains appeared to be infected and to be lysed equally (Figure 2E, t= 5, 9 hrs; Movie S4). However, during the constriction phase, when the bacteria re-grew into the plaque zone, WT cells outcompeted the *ΔsigX* cells, as signified by the dominant colonization of the mCherry-labeled WT cells at the plaque rim (Figure 2E, t=12, 16 hrs; Figure S3A; Movie S4). When GFP- and mCherry-labeled WT cells were mixed as a control, both strains were evenly distributed at the plaque edge even 16 hours post infection (Figure S3A). Consistent with these results, fusion of the *sigX* promoter to *gfp* specified that cells located at the plaque rim produced GFP chiefly during the constriction phase (Figure 2F). In addition, cell infection at 48°C, a temperature shown to activate *sigX* expression (Huang et al., 1997), led to a significant reduction in plaque size in a *sigX*-dependent manner, without any measurable impact on growth kinetics (Figure S3B-S3C). We conclude that *ΔsigX* cells are deficient in inducing a defense mechanism that enables bacteria to tolerate the presence of phages and invade into the plaque zone.

### SigX is activated in non-infected bacteria following infection of their neighbors

The role of SigX in inducing the tolerance response was further substantiated by monitoring the production and localization of SigX during infection at the cellular level. In the absence of phages, SigX fused to GFP (P*sigX-sigX-gfp*) mainly localized onto the membrane, frequently forming foci assemblies in proximity to the cell circumference and at septal positions (Figure 3A). This localization pattern is consistent with previous reports showing that SigX is sequestered to the plasma membrane by its anti-sigma factor as a way to halt its action (Ho and Ellermeier, 2012). To assay SigX activity in uninfected bacteria, we added SPP1 phage to mCherry-labeled WT bacteria mixed with phage resistant bacteria, lacking the SPP1 receptor (*ΔyueB*) and harboring *sigX-gfp*. Time-lapse microscopy revealed repositioning of SigX-GFP from membrane and foci locations to massive nucleoid deployment in the resistant bacteria (Figure 3B; t=35 min), indicating a switch from an inactive to an active mode. Noticeably, this shift in localization occurred prior to lysis of nearby infected sensitive bacteria. SigX-GFP level was dropped and its localization into foci was largely restored in the resistant bacteria 95 min post infection (Figure 3B), presumably corresponding to conclusion of the damage sensing response. Taken together, SigX appears to be activated in phage resistant bacteria upon sensing a damage signal from nearby infected sensitive cells. Of note, infecting *sigX-gfp* sensitive cells with SPP1 showed that SigX-GFP largely displaces its position form the membrane to the nucleoid in the course of infection (Figure S3D; t=30 min), denoting that also infected cells activate the SigX response.

**Figure 3.**
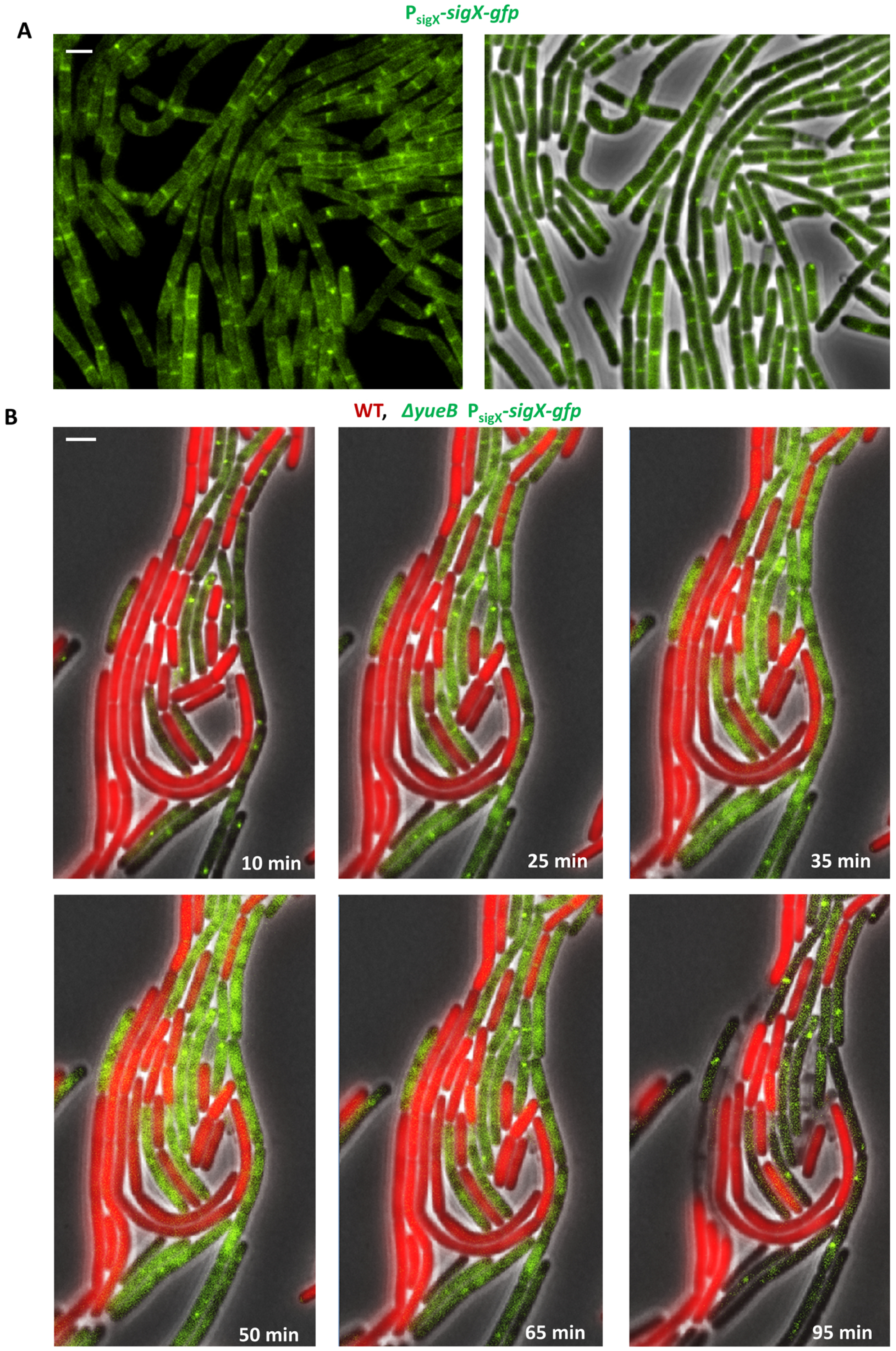
SigX is activated in non-infected cells upon infection of their neighbors. **(A)** ET26 (P_sigX_*-sigX-gfp*) cells were placed on an agarose pad and followed by time lapse fluorescence microscopy. Shown are signal from SigX-GFP (green) (left panel), and an overlay image of phase contrast (grey) and signal from SigX-GFP (green) (right panel), captured at the indicated time points. **(B)** BDR2637 (P_veg_*-mCherry*) (WT, red) and ET261 (*ΔyueB,* P_sigX_*-sigX-gfp*) (green) cells were mixed, infected with SPP1, placed on an agarose pad and followed by time lapse fluorescence microscopy. Shown are overlay images from mCherry (red), SigX-GFP (green), and phase contrast (grey), captured at the indicated time points. Scale bars 1 μm.

### Expression of SigX protects from phage infection

The impact of SigX on phage infection was further explored by constructing bacteria artificially expressing SigX under an IPTG-inducible promoter. Remarkably, expressing *sigX* prior to phage addition markedly attenuated both SPP1 and Phi29 infections, with the cells being capable of extending the infection process (Figure 4A). Next, mCherry-labeled cells, over-expressing SigX (P_IPTG_*-sigX*), were incubated with non-labeled WT cells, and the mixture was infected with SPP1-*_lysin-yfp_*. Consistent with the above observations, WT cells were rapidly infected and lysed, while cells over-expressing SigX appeared to be infected at slower kinetics and to a lesser extent (Figure 4B-4C), a phenomenon that was also observed during infection with Phi29 (Figure S4A).

**Figure 4.**
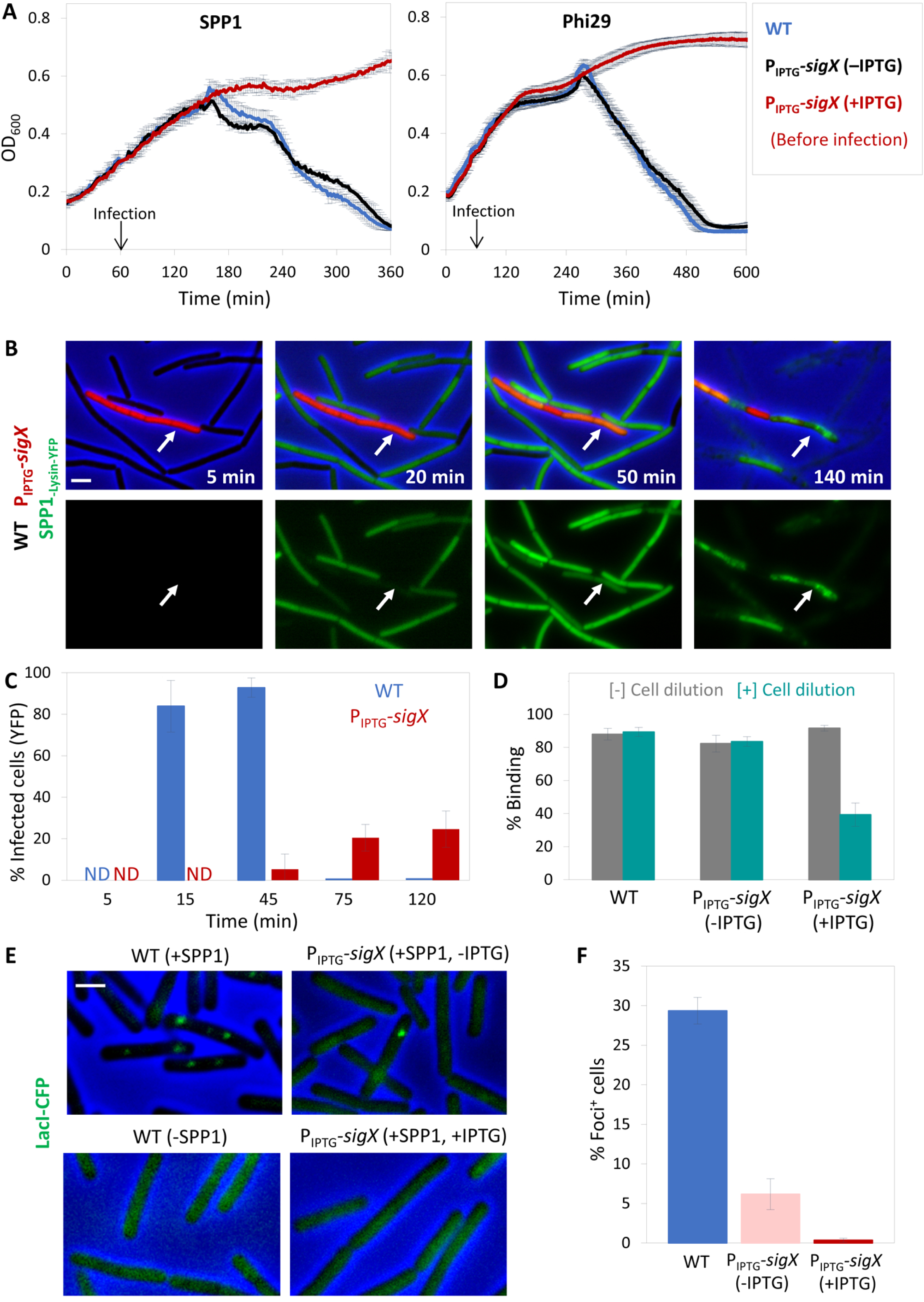
SigX expression confers phage tolerance. **(A)** PY79 (WT) and ET28 (P_IPTG_*-sigX*) cells were infected with SPP1 or Phi29 (t=60 min) at 1:20 (phages:bacteria) MOI, and OD_600_ was followed at 2 min intervals. IPTG was added 30 min before infection (t=30 min). Shown is a representative experiment out of 3 biological repeats, and the average values and SD of 4 technical repeats. **(B)** ET29 (P_veg_*-mCherry*, P_IPTG_*-sigX*) (red) cells were grown in the presence of IPTG and mixed with PY79 (WT) cells. The mixture was infected with SPP1-*_lysin-yfp_* 1:1 (phages:bacteria) MOI, placed on an IPTG-containing agarose pad, and followed by time lapse fluorescence microscopy. Shown are overlay images of phase contrast (blue), signal from mCherry labeled cells (red), and signal from Lysin-_SPP1_-YFP (green), captured at the indicated time points (upper panels). Corresponding signal from Lysin-_SPP1_-YFP (green) is shown separately (lower panels). Arrows highlight the delayed infection of ET29 cells. Scale bar 1 μm. **(C)** Quantification of the experiment described in (B). Shown is the percentage of phage infected PY79 (WT) and ET29 (P_veg_*-mCherry*, P_IPTG_*-sigX*) cells at the indicated time points, scored by the Lysin-_SPP1_-YFP signal, with average values and SD (n ≥ 200 cells for each time point). Of note, the majority of WT cells were lysed at t=75 min post infection. **(D)** PY79 (WT) and ET28 (P_IPTG_*-sigX*) cells, grown in the presence or absence of IPTG, were infected with SPP1 (1:1 MOI) for 10 min. Next, phage adsorption was monitored before and after cell dilution (x100 fold). Percentage of phage adsorption was calculated as follows: (P_0_-P_1_)x100/P_0_, where P_0_ is the initial phage input in the lysate (PFU/ml), and P_1_ is the titer of free phages (PFU/ml) 10 min after cell infection. Shown are average values and SD of a representative experiment out of 3 independent experiments. **(E)** OF83 (P_pen_*-lacIΔ11-cfp*) (WT) and ET40 (P_pen_*-lacIΔ11-cfp,* P_IPTG_*-sigX*) cells, grown in the presence or absence of IPTG, were infected with SPP1-*_delX110lacO64_*. The formation of LacI-CFP foci on phage DNA was monitored 10 min post infection. OF83 cells without infection were used for comparison. Shown are overlay images of phase contrast (blue) and signal from LacI-CFP (green). Scale bar 1 µm. **(F)** Quantification of the experiment described in (E). Shown is the percentage of LacI-CFP foci 10 min post infection of OF83 and ET40 cells by SPP1, with average values and SD (n ≥ 850 cells for each population).

To define the specific stage at which phage infection was interrupted by SigX activity, we followed the adsorption of phages to cells over-expressing *sigX*. A standard adsorption assay yielded no significant difference in SPP1 adsorption rate between WT and SigX expressing cells (Figure 4D). Nonetheless, since SPP1 phage exhibits two modes of cell surface binding, reversible and irreversible (Baptista et al., 2008), it was still conceivable that the irreversible mode of binding was impaired. To inspect this possibility, cells were diluted after an initial phage adsorption period to enable reversibly adsorbed phages to detach from the host (Baptista et al., 2008). Indeed, a large fraction of phages, adsorbed to *sigX* over-expressing cells, were liberated after dilution, whereas no significant release of WT-attached phages was detected (Figure 4D), indicating that SigX expression delays SPP1 irreversible binding. To corroborate this finding, we investigated whether phage DNA injection is consequently delayed by induced expression of SigX. To monitor phage DNA injection, we utilized SPP1_-*delX110lacO64*_ phage, which contains 64 repeats of *lacO* (Jakutyte et al., 2012), and infected bacteria chromosomally expressing *lacI-cfp*. The presence of phage DNA within the host cytoplasm was visualized subsequent to injection by the formation of LacI-CFP foci (Fernandes et al., 2016; Tzipilevich et al., 2017). Infection of WT cells resulted in foci appearance within 10 minutes after infection, while no foci were observed in *sigX* over-expressing cells at the same time point (Figure 4E-4F), indicating a delay in phage DNA penetration into the latter cell population. Consistent with this possibility, a significant decrease in SPP1-mediated plasmid transduction rate into SigX producing cells was monitored (Figure S4B). Thus, we surmise that *sigX* expression interferes with SPP1 irreversible binding and consequently delays phage DNA injection.

### The *dlt* operon mediates the SigX tolerance response to phage infection

To identify the gene(s) required for SigX-mediated phage tolerance, we mutated known genes in the SigX regulon (Huang and Helmann, 1998), and tested their impact on the phage protection phenotypes. Intriguingly, disruption of the *dlt* operon (Δ*dltA*), largely countered the tolerance to phage infection conferred by SigX over-expression (Figure 5A and Figure S5A). The *dlt* operon encodes an enzymatic pathway that is known to ligate D-alanine moieties to TA polymers (Perego et al., 1995), the major phage surface attachment components. Examination of infected *ΔdltA* mutant cells harboring inducible *sigX* by fluorescence microscopy further substantiated that they were lysed with kinetics similar to that of WT cells in the presence of the inducer (Figure S5B). Furthermore, deletion of *dltA* restored the capacity of SPP1 to bind irreversibly to the surface of *sigX* over-expressing cells, and consistently increased the level of SPP1 DNA injection into these cells, as detected by transduction assay (Figure 5B-5C).

**Figure 5.**
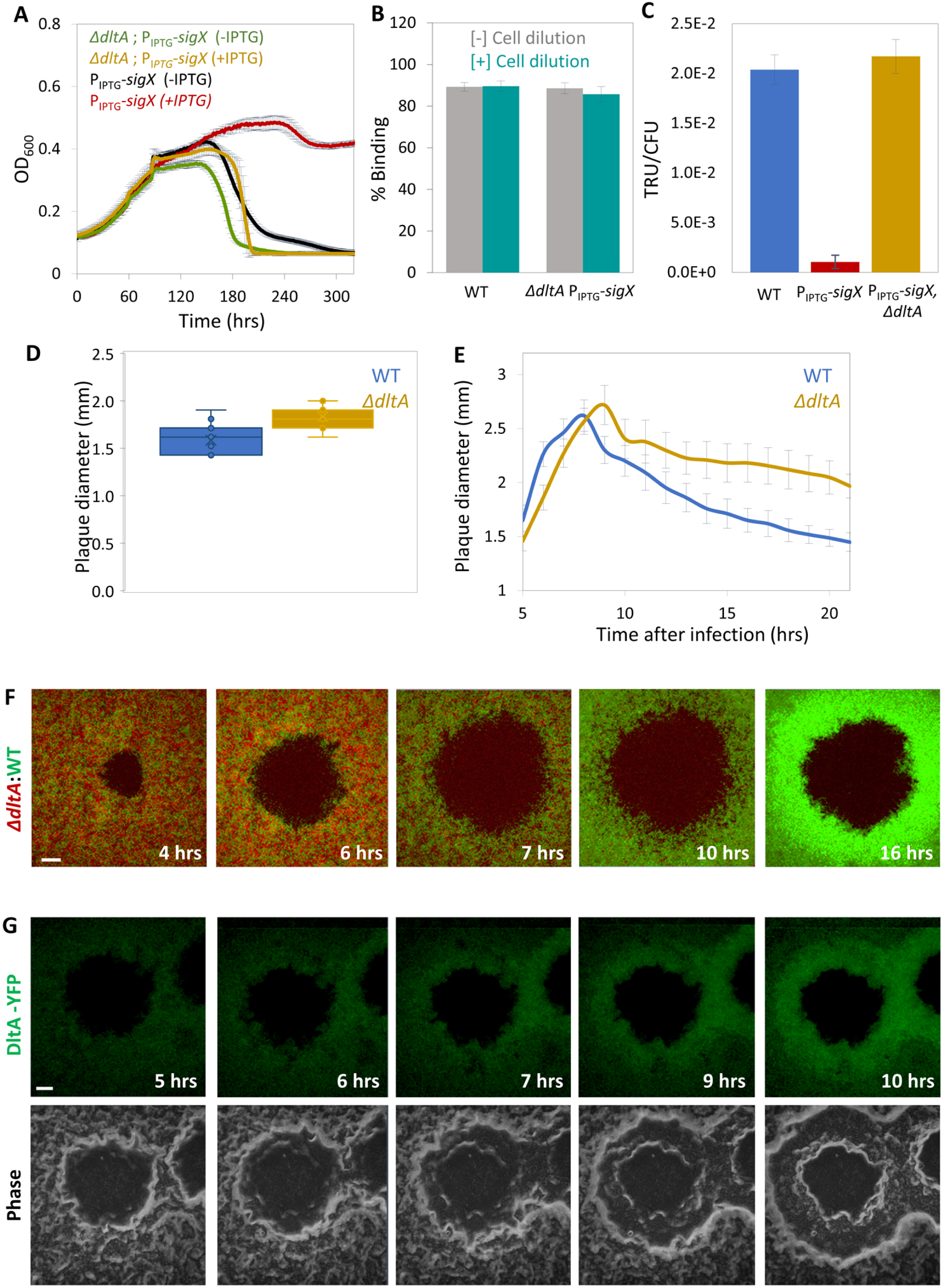
The *dlt* operon mediates the SigX-induced phage tolerance response. **(A)** ET28 (P_IPTG_*-sigX*) and ET42 (Δ*dltA*, P_IPTG_*-sigX*) cells, grown in the presence or absence of IPTG, were infected with SPP1 at 1:20 (phages:bacteria) MOI, and OD_600_ was followed at 2 min intervals. Shown is a representative experiment out of 6 biological repeats, with the average values and SD of 8 technical repeats. **(B)** PY79 (WT) and ET42 (*ΔdltA*, P_IPTG_*-sigX*) cells, grown in the presence of IPTG, were infected with SPP1 (1:1 MOI) for 10 min. Next, phage adsorption was monitored before and after cell dilution (x100 fold). Percentage of phage adsorption was calculated as follows: (P_0_-P_1_) x100/P_0_, where P_0_ is the initial phage input in the lysate (PFU/ml), and P_1_ is the titer of free phages (PFU/ml) 10 min after cell infection. Shown are average values and SD of 5 biological repeats. **(C)** PY79 (WT), ET28 (P_IPTG_*-sigX*), and ET42 (*ΔdltA,* P_IPTG_*-sigX*), grown in the presence of IPTG, were transduced with SPP1-pBT163 lysate, and the number of transductants was monitored by plating the cells on selective plates. Transduction unit (TRU) was calculated as the number of transductant colonies /total CFU. Shown are average values and SD of 3 biological repeats. **(D)** PY79 (WT) and ET41 (*ΔdltA*) cells were infected with SPP1 and plaque diameter was monitored after 20 hrs of incubation. Shown is plaque diameter distributions for each strain (n ≥ 40). **(E)** Plaque formation dynamic of SPP1 was monitored by automated scanning (Levin-Reisman et al., 2010) on a lawn of infected PY79 (WT) or ET41 (*ΔdltA*) cells grown on an MB agar plate. Shown are average values and SD from kinetics random plaques for each strain (n ≥ 10). **(F)** AR16 (P_rrnE_*-gfp*) (WT, green) and ET411 (P_veg_*-mCherry, ΔdltA*) (*ΔdltA*, red) cells were mixed, infected with low concentrations (10^-8^ PFU/ml) of SPP1, placed on an agarose pad, and plaque formation was followed by time lapse confocal microscopy. Shown are overlay images from GFP (green) and mCherry (red) signals of the bacterial lawn captured at the indicated time points (hrs). The plaque is seen as a hole formed on the bacterial lawn. Scale bar 150 µm. **(G)** ET43 (*dltA-yfp)* cells were infected with low concentrations (10^-8^ PFU/ml) of SPP1, placed on an agarose pad, and plaque formation was followed by time lapse confocal microscopy. Shown are fluoresce from DltA-YFP signal (upper panels) and corresponding phase contrast images (lower panels), captured at the indicated time points (hrs). The plaque is seen as a hole formed on the bacterial lawn. Scale bar 100 µm.

To substantiate the role of the *dlt* operon in the phage tolerance response, we examined the phenotype of *ΔdltA* in an otherwise WT background. *ΔdltA* cells exhibited plaques larger than that of the WT, along with prominent deficiency in plaque constriction phase (Figure 5D-5E; Figure S5C), similar phenotypes to those observed for the *ΔsigX* cells (Figure 2A-2B). Next, we mixed differentially labeled *ΔdltA* and WT cells, and followed plaque formation dynamics by time lapse confocal microscopy. While the two populations were evenly distributed at early stages of plaque generation, WT cells were manifestly dominating the plaque rim during constriction (Figure 5F), indicating clear deficiency of the mutant cells in opposing infection. Monitoring DltA expression by following DltA-YFP showed an enrichment of the fusion protein preferentially at the edge of the constricting plaque (Figure 5G), supporting a role in mediating this process. In sum, the majority of the phage protection phenotypes conferenced by SigX can be assigned to the *dlt* operon, encoding enzymes that modify the TA surface polymers.

## Discussion

Plaque signifies phage attack, providing a visible mark to trace and monitor infection. However, mechanisms elucidating plaque expansion and confinement are hardly described (Abedon and Yin, 2009). By following the dynamics of plaque formation on lawns of the soil bacterium *B. subtilis,* we discovered that plaque development includes a phase of constriction, typified by bacterial growth into the plaque zone, counteracting plaque expansion. Examination of bacteria located at the plaque rim subsequent to constriction revealed that they are not genetically resistant to phages, but instead elicit a temporary phage tolerance response, activated by the stress-induced RNA polymerase sigma factor σ^X^. We further uncovered that the impact of SigX on tolerance is mostly attributed to the action of the *dlt* operon, encoding for WTA modifying enzymes, thereby altering the polymer properties. Based on our results, we propose that uninfected bacteria can sense infection of their neighbors, and in turn trigger a tolerance response, modifying their phage attachment surface components to antagonize phage penetration (Figure 6). As such, the cells at the plaque rim form a protective barrier that locally constrains plaque spread, shielding the non-infected population.

**Figure 6.**
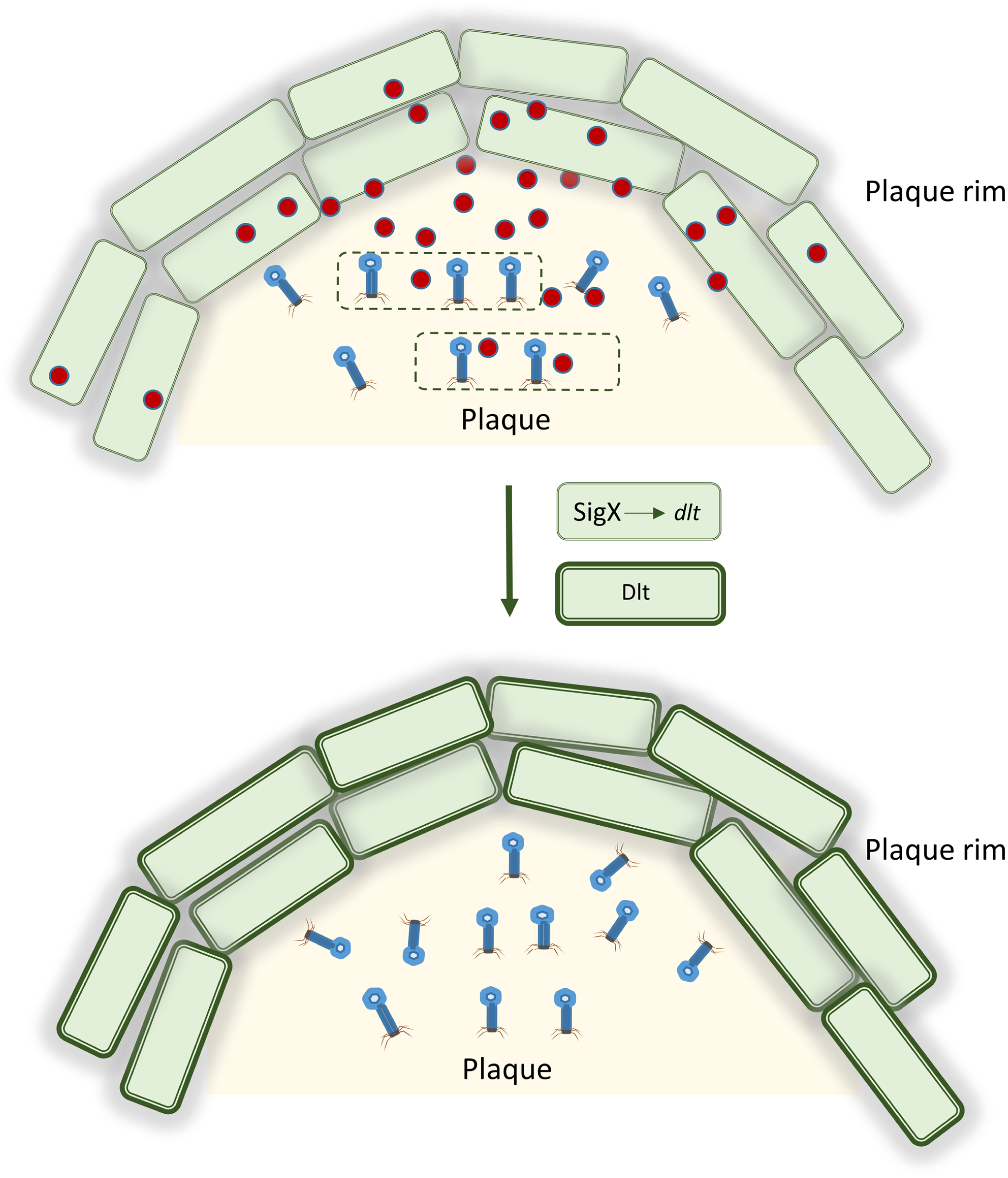
**A model for eliciting phage tolerance response by uninfected bacteria** Upon phage infection bacteria (dashed-line cells) generate a yet unidentified “danger signal” (red circles) that is received by uninfected neighboring cells located at the plaque rim. In turn, the latter cells activate SigX that induces the transcription of the *dlt* operon. The produced Dlt enzymes modulate the phage receptor WTA polymers to reduce phage binding, providing temporary protection against phage attack and limiting plaque spread.

This scenario resembles the eukaryotic innate immune response, with the hallmark being interferon released by infected cells and received by neighbors to activate an anti-viral response (Isaacs and Lindenmann, 1957; McNab et al., 2015). In addition, endogenous danger signals, such as extracellular ATP and DNA, released by infected eukaryotic cells, stimulate innate immunity (Gallucci and Matzinger, 2001), a program that could be applicable for activating the phage tolerance response. Importantly, damaged-self recognition stimulated by similar factors exists in plants and even in algae and fungi (Heil and Land, 2014). Interestingly, tt has been shown that bacteria possess the cyclic GMP–AMP synthase (cGAS)–STING pathway, a central component of the mammalian innate immune system, as part of an anti-phage defense system (Cohen et al., 2019; Morehouse et al., 2020).

Our data indicate that bacteria acquire phage tolerance chiefly by remodeling their cell surface, decorating WTA polymers with D-alanine residues (Brown et al., 2013). D-alanylation of TA polymers is executed by a series of chemical reactions performed by the Dlt enzymes (Ma et al., 2018; Percy and Grundling, 2014; Perego et al., 1995). It has been shown that TA D-alanylation contributes to resistance against cationic antimicrobial peptides and lysozyme (Kingston et al., 2013; Kovacs et al., 2006), presumably due to changes in the cell surface electric charge or increase in peptidoglycan density (Percy and Grundling, 2014; Saar-Dover et al., 2012). D-alanylation of TA was also shown to enhance host cell adhesion and virulence of Gram positive pathogens such as *Staphylococcus aureus* and *B. anthracis* (Percy and Grundling, 2014; Simanski et al., 2013). Since TA glycosylation is known to serve as a pervasive phage binding molecule (Habusha et al., 2019; Young, 1967), it is plausible that WTA-D-alanylation masks WTA glycosylated sites, and/or interferes with bacteriophage access to the membrane.

The induction of the phage tolerance response was found to be mediated by SigX, belonging to the ECF family that monitors cell wall integrity. SigX is recruited to the membrane by its cognate anti-sigma factor and is liberated to the nucleoid in response to envelope stress to transcribe downstream target genes, with one of the most prominent being the *dlt* operon (Cao and Helmann, 2004; Helmann, 2016). A *sigX* mutant strain was shown to be sensitive to heat and oxidative stress, and to be susceptible to cationic antimicrobial peptide (Cao and Helmann, 2004; Huang et al., 1997). Indeed, we found that elevated growth temperature indirectly protects bacteria from phage attack in a SigX-dependent manner, hinting that cross activation of ECF enables bacteria to simultaneously resist multiple stress conditions, similarly to SOS response (Gottesman, 2019; Storz, 2016). Still, the nature of the activating signals and how they are transduced to release the membrane-attached SigX, have yet to be elucidated. Since ECFs are widespread among bacteria (Helmann, 2002), it is tempting to assume that, similarly to *B. subtilis*, many species have the capacity to execute such a defense strategy, following infection of nearby bacteria.

Bacteria have evolved numerous remarkable phage resistance strategies to counteract infection, including restriction-modification systems, abortive infection, CRISPR-Cas immunity (Labrie et al., 2010; Salmond and Fineran, 2015), and a plethora of recently identified additional exciting systems [e.g. (Cohen et al., 2019; Doron et al., 2018; Gao et al., 2020; Makarova et al., 2011)]. Still, relatively little is known about the dynamics of the activation of these systems within populations. For instance, it is not entirely understood how activation of the CRISPR-Cas system is prompted. There is evidence for constitutive activity of *Cas* genes in some bacteria, and for quorum sensing mediated transcription of *Cas* genes in others (Hampton et al., 2020; Patterson et al., 2017). Consistently, proteomic analysis revealed that phage infection elevates Cas production in *Streptococcus thermophilus* (Young et al., 2012). Knowledge concerning population dynamics of other phage defense systems is even more fragmentary. Here we uncovered a general phage tolerance mechanism that provides only temporary protection by phenotypic modulation of the bacterial cell surface. To the best of our knowledge, this is the first detailed characterization of a phage defense system in space and time.

## Acknowledgements

We thank I. Rosenshine and A. Rouvinski (Hebrew University, Israel), and members of the Ben-Yehuda laboratory for valuable discussions. We are grateful to P. Tavers (Gif-sur-Yvette, France), J. Helmann (Cornell University, USA) and D. Rudner (Harvard University, USA) for providing strains. This work was supported by the NSF/BSF-United States-Israel Binational Science Foundation (2017672), and the ERC Synergy grant (810186) awarded to S. B-Y.

## Supplementary Figures

**Figure S1.**
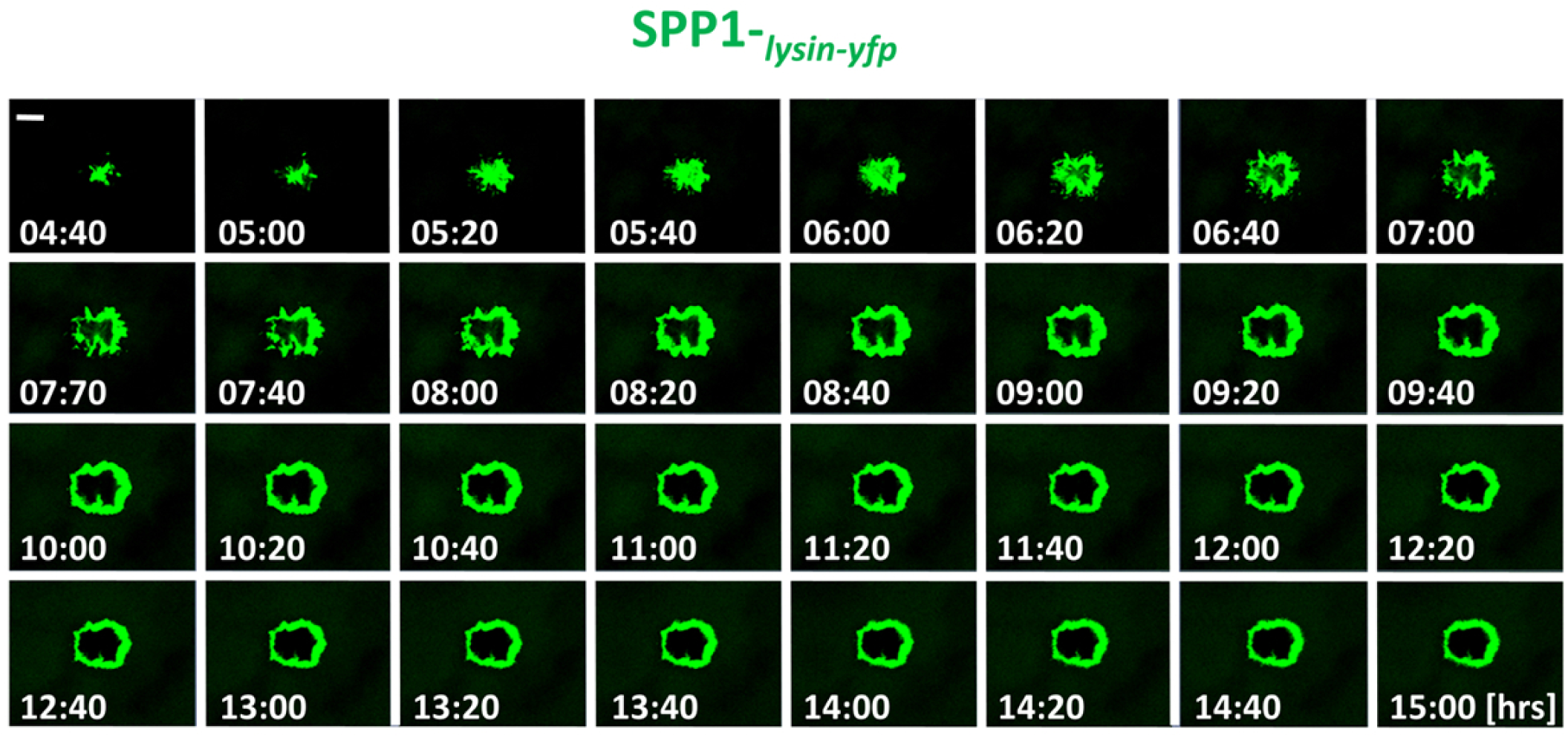
Phages can be visualized at the plaque periphery. PY79 (WT) cells were infected with low concentrations (10^-8^ PFU/ml) of SPP1-*_lysin-yfp_*, placed on an agarose pad and plaque formation was followed by time lapse confocal microscopy. Shown is the signal from Lysin-_SPP1_-YFP (green), captured at the indicated time points (hrs). Scale bar 50 µm. Corresponds to Movie S3.

**Figure S2.**
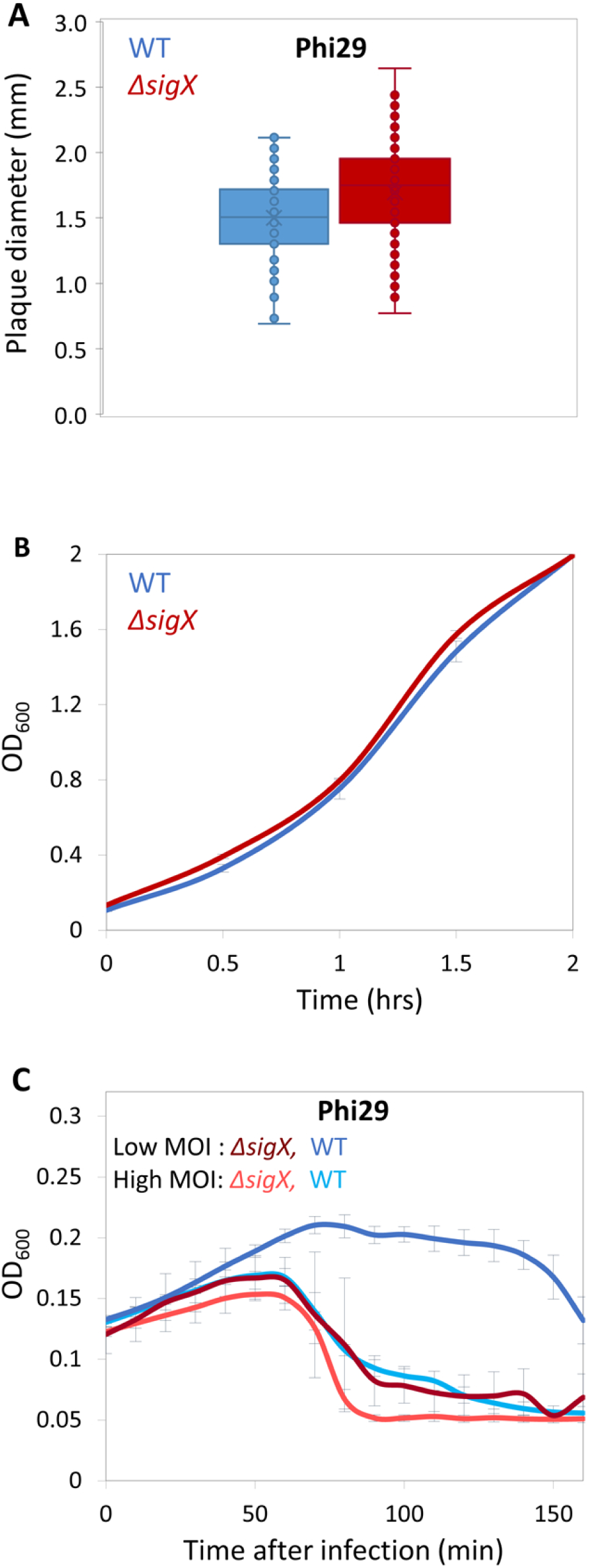
*ΔsigX* cells are highly sensitive to Phi29 infection. **(A)** PY79 (WT) and ET19 (*ΔsigX*) cells were infected with Phi29 and plaque diameter was monitored after 20 hrs of incubation. Shown is plaque diameter distribution for each strain (n ≥ 125). **(B)** PY79 (WT) and ET19 (*ΔsigX*) strains were grown in LB liquid medium and OD_600_ monitored. Shown are average values and SD of 3 biological repeats. **(C)** PY79 (WT) and ET19 (*ΔsigX*) cells were infected with Phi29 at either high (phages:bacteria 1:1) or low (phages:bacteria 1:20) MOI, and OD_600_ was followed at 5 min intervals. Shown is a representative experiment out of 3 biological repeats, and average values and SD of 8 technical repeats.

**Figure S3.**
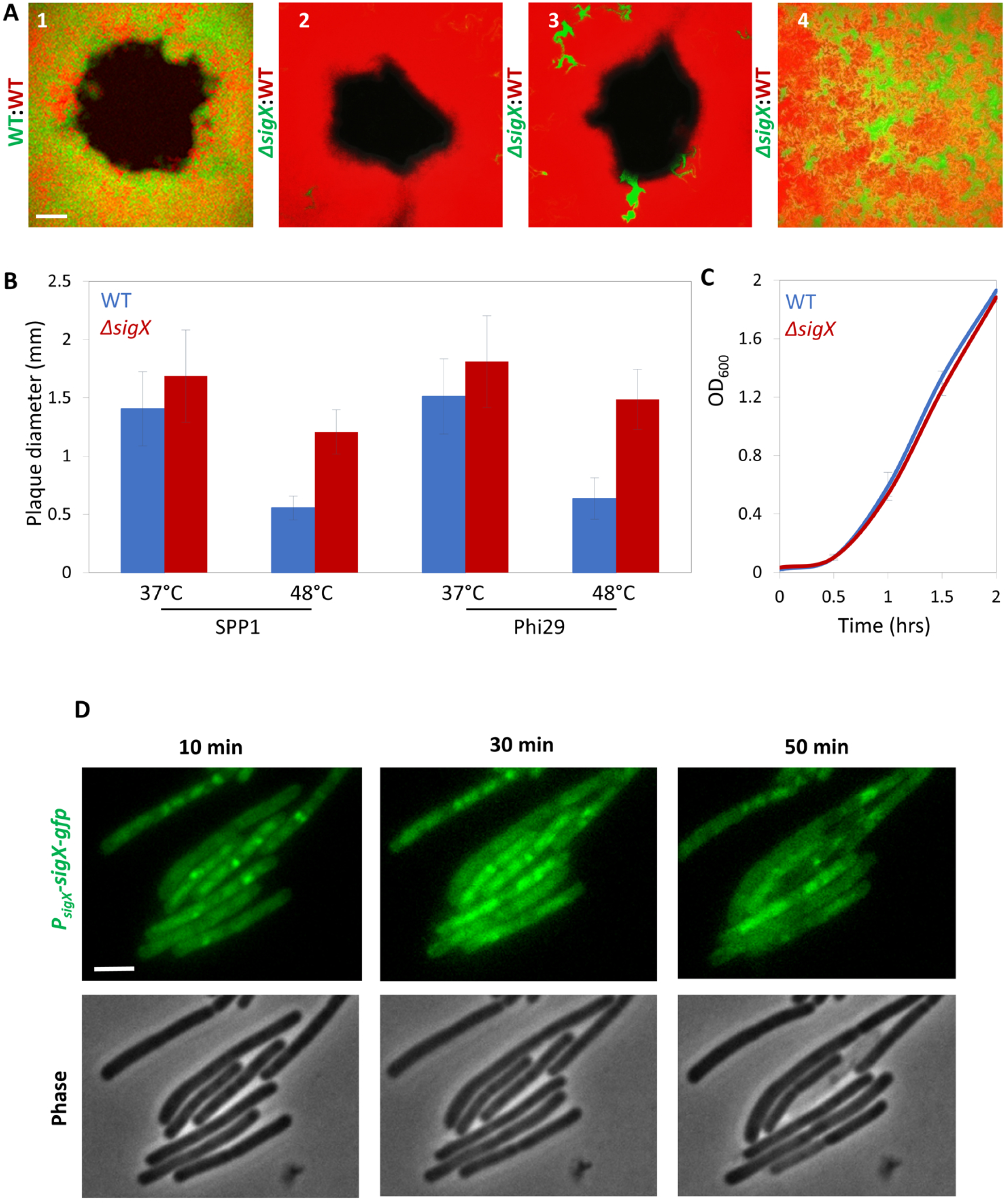
SigX activation mediates phage tolerance. **(A)** BDR2637 (P_veg_*-mCherry*) (red) cells were mixed with AR16 (P_rrnE_*-gfp*) (green) **(1)** or with ET191 (*ΔsigX,* P_rrnE_*-gfp*) (green) **(2-4)** cells. The mixtures were infected with low concentrations (10^-8^ PFU/ml) of SPP1, placed on an agarose pad and plaque formation was followed by time lapse confocal microscopy. Shown are overlay images of mCherry (red) and GFP (green) signals captured 16 hrs post infection. (1-3) show plaque regions whereas (4) shows a region remote from any visible plaque site. Scale bar 100 µm. **(B)** PY79 (WT) and ET19 (*ΔsigX*) cells were infected with SPP1 or Phi29 and plates were incubated either at 37°C or 48°C. Plaque diameter was monitored after 20 hrs of incubation (n ≥ 60). Shown are average values and SD of 3 independent repeats. **(C)** PY79 and ET19 were grown in liquid LB medium at 48°C and OD_600_ monitored. Shown are average values and SD of 3 biological repeats. **(D)** ET26 (P_sigX_*-sigX-gfp*) cells were infected with SPP1, placed on an agarose pad and followed by time lapse fluorescence microscopy. Shown are signal from SigX-GFP (top panels), and phase contrast SigX-GFP images (bottom panels), captured at the indicated time points post infection. Scale bar 1 µm.

**Figure S4.**
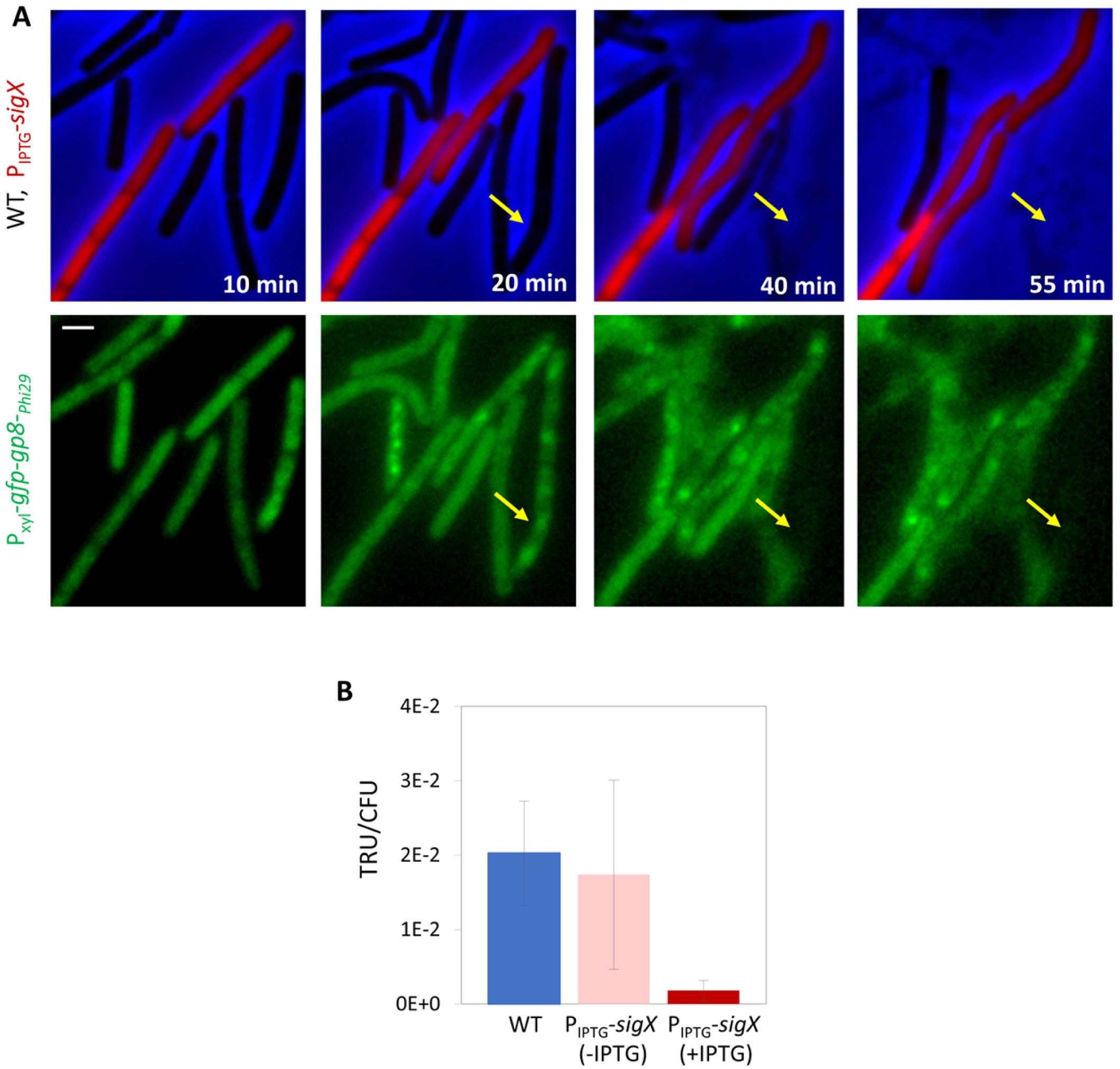
SigX over-expression interferes with phage infection. **(A)** ET9 (P_xyl_*-gfp-gp8*-*_Phi29_*) (WT) and ET44 (P_veg_*-mCherry*, P_IPTG_*-sigX,* P_xyl_*-gfp-gp8_-Phi29_*) (P_IPTG_*-sigX,* red) cells were grown in the presence of IPTG and xylose, mixed, and infected with Phi29. The mixture was placed on an IPTG and xylose-containing agarose pad and followed by time lapse fluorescence microscopy. Gp8-_Phi29_ is the major Phi29 capsid protein that localizes into discrete foci during Phi29 infection (Tzipilevich et al., 2017). Shown are overlay images of phase contrast (blue) and signal from mCherry (red) (upper panels), and signal from GFP-Gp8-_Phi29_ (green) (lower panels), captured at the indicated time points. Arrows highlight GFP-Gp8-_Phi29_ foci appearance in ET9 cells. Scale bar 1 µm. **(B)** PY79 (WT) and ET28 (P_IPTG_*-sigX*), grown in the presence or absence of IPTG, were transduced with SPP1-pBT163 lysate, and the number of transductants was monitored by plating the cells on corresponding selective plates. Transduction unit (TRU) was calculated as the number of transductant colonies /total CFU. Shown are average values and SD of 3 biological replicates.

**Figure S5.**
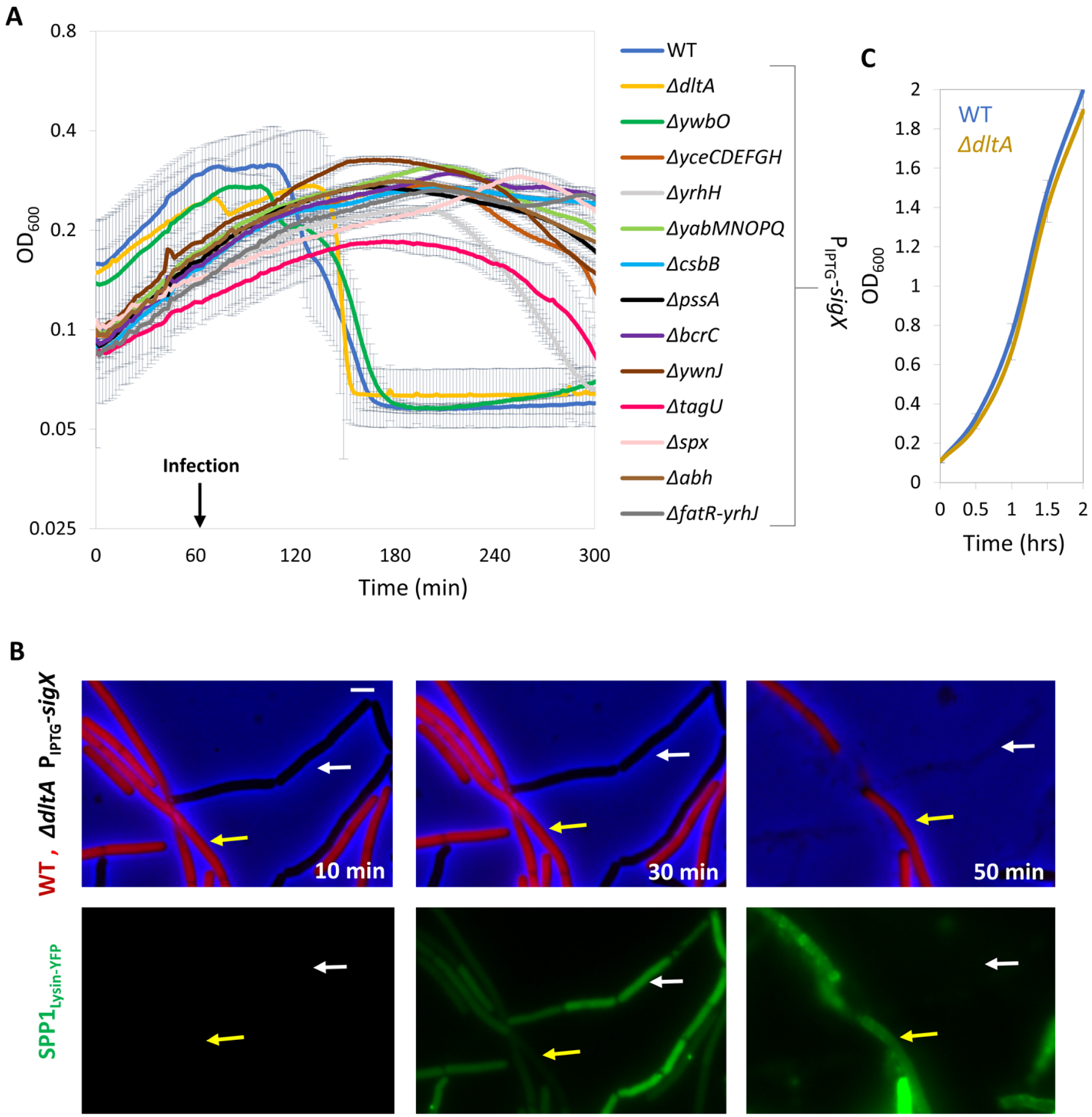
SigX impact on phage tolerance is Dlt-mediated. **(A)** Bacterial strains harboring P_IPTG_*-sigX* as well as the indicated gene deletions were grown in the presence of IPTG. At t=60 min, cells were infected with SPP1 at low (phages:bacteria 1:20) MOI, and OD_600_ was followed at 2 min intervals. PY79 (WT) was infected in parallel for comparison. *dltA* and *ywbO* mutations diminished the resistance gained by *sigX* expression. Shown is a representative experiment out of 3 independent biological repeats, with average values and SD of 3 technical repeats. **(B)** ET42 (*ΔdltA,* P_IPTG_*-sigX*) cells were grown in the presence of IPTG and mixed with BDR2637 (P_veg_*-mCherry*) (WT, red) cells. The mixture was infected with SPP1-*_lysin-yfp_*, placed on an IPTG-containing agarose pad, and followed by time lapse fluorescence microscopy. Shown are overlay images of phase contrast (blue) and signal from mCherry labeled cells (red) (upper panels), and the corresponding signal from Lysin-_SPP1_-YFP (green) (lower panels), captured at the indicated time points. Yellow arrows denote infected WT cells whereas white arrows highlight infected ET42 cells that lysed rapidly. Scale bar 1 μm. **(C)** PY79 (WT) and ET41 (*ΔdltA*) cells were grown in LB liquid medium and OD_600_ was followed. Shown are average values and SD of 3 biological repeats.

## Supplementary Movie Legend

**Movie S1. Plaque expansion is followed by a phase of plaque constriction**

BDR2637 (P*_veg_-mCherry*) cells were infected with low concentrations (10^-8^ PFU/ml) of SPP1, placed on an agarose pad and plaque formation was followed for 15 hrs by time lapse confocal microscopy. Shown are overlay images, captured at 20 min intervals, from mCherry signal (red) and phase contrast (grey) of the bacterial lawn. The plaque is seen as a hole formed on the bacterial lawn. Corresponds to Figure 1A.

**Movie S2. Following plaque dynamics on agar plates by automated scanning**

Plaque formation was monitored by automated scanning (Levin-Reisman et al., 2010) on a lawn of PY79 (WT) cells grown on an MB agar plate. Images were captured at 15 min intervals in the course of 19 hrs. Corresponds to Figure 1B.

**Movie S3. Phages can be visualized at the plaque periphery**

PY79 (WT) cells were infected with low concentrations (10^-8^ PFU/ml) of SPP1-*_lysin-yfp_*, placed on an agarose pad and plaque formation was followed by time lapse confocal microscopy. Shown are overlay images from Lysin-_SPP1_-YFP (green) and phase contrast (grey) of the bacterial lawn captured at 20 min intervals in the course of 15 hrs. Corresponds to Figure S1.

**Movie S4. ΔsigX cells are deficient in plaque constriction**

BDR2637 (P_veg_*-mCherry*) (WT, red) and ET191 (P_rrnE_*-gfp*, *ΔsigX*) (*ΔsigX*, green) cells were mixed, infected with low concentrations (10^-8^ PFU/ml) of SPP1, placed on an agarose pad and plaque formation was followed by time lapse confocal microscopy. Shown are overlay images from mCherry (red) and GFP (green) signal of the bacterial lawn captured at 20 min intervals in the course of 16 hrs. The plaque is seen as a hole formed on the bacterial lawn. Corresponds to Figure 2E.

## METHODS

### Strains and plasmids

*B. subtilis* strains were derivatives of the wild-type PY79 (Youngman et al., 1984). All bacterial strains and phages are listed in Table S1. Plasmid constructions were performed in *E. coli* DH5α using standard methods and are listed in Table S1. All primers used in this study are listed in Table S2.

### General growth conditions

Bacterial cultures were inoculated at OD_600_ 0.05 from an overnight culture and growth was carried out at 37°C in LB medium supplemented with 5 mM MgCl_2_ and 0.5 mM MnCl_2_ (MB) (Harwood and Cutting, 1990). For induction of all P*_hyper-spank_* controlled genes were induced with Isopropyl-β-D-thiogalactopyranoside (IPTG) at concentration of 0.5 mM.

### Phage lysis and transduction

Phage lysate was prepared by adding approximately 10^9^ phages to mid-log cells grown in MB until the culture was completely cleared. Next, the lysate was filtered through 0.45 µm or 0.22 µm Millipore filter. For lysis dynamic experiments, phages were added to mid-log growing cells at the indicated MOI, and OD_600_ was monitored. For sigma X induction experiments, cells were grown for 30 min in the absence of inducer. Subsequently, inducer was added and cells were grown for additional 30 min. Phages were added after overall growth of 60 min and OD_600_ followed by Spark 10M (Tecan) multiwell fluorometer set at 37°C with a constant shaking. In the experiments by which SigX activity was tested during the infection cycle the inducer and the phages were added simultaneously at 30 min.

For transduction experiments, lysates were prepared from a given donor strain as describe above. All lysates for transduction were treated with DNase I (Sigma Aldrich) 200 ng/ml for 20 min at RT. Recipient strains were grown to 1 OD_600_, and cells (1 ml) were mixed with 100 µl of lysate and 9 ml of MB, and incubated at 37°C without shaking for 20 min. Subsequently, cells were centrifuged and spread on selective plates supplemented with 10 mM sodium citrate (Clokie and Kropinski, 2009). For burst size measurement 1:10^4^ phages:bacteria were added to a mid-log culture, 30 minutes later the culture was lysed with chloroform and the number of phages in the culture was measured.

### Phage attachment assay

Indicated bacterial strains were grown in a liquid culture to 0.6 OD_600_, and phage adsorption to the cells was measured at 10 min post infection, by titrating the free phages present in the supernatant as previously described (Ellis and Delbruck, 1939). In brief, logarithmic cells were grown in MB at 37°C till 0.8 OD_600_, then 15 mM CaCl_2_ and 50 µg chloramphenicol/ml were added to the medium and cells were incubated for 10 min. Next, cells were infected with phages (10^7^ PFU/ml), and samples (0.5 ml) were collected at 10 min post infection, centrifuged for 1 min, and 50 µl of the supernatant was diluted, plated, and PFU/ml was determined. To measure reversible attachment, 10 min post infection, cells were diluted 100 fold in fresh MB, incubated for 2 min in 37°C and free phages present in the supernatant were plated and PFU/ml was determined.

### Plaque size determination

For final plaque size determination, bacteria from mid-log culture were infected with low concentration of phages (MOI=10^-6^) and incubated on plates at 37°C or 48°C for 20 hrs. Next, plates were photographed and plaque size was determined by measuring the size of random plaques. Image processing was performed using MetaMorph 7.4 software (Molecular Devices). For analyzing plaque growth dynamics on agarose plates, bacteria were infected as describe above, and plates were incubated at 37°C on top of a scanner and covered with a dark paper cover. An automated scanning program (Levin-Reisman et al., 2010) was utilized for time lapse imaging of the plates. Plaque size was determined by measuring the size of random plaques at intervals of 1 hr. Image processing was performed using MetaMorph 7.4 software (Molecular Devices).

### Fluorescence microscopy

For fluorescence microscopy, bacterial cells (0.5 ml, OD_600_ 0.5) were centrifuged and suspended in 50 μl of MB. For time lapse microscopy, bacteria were placed over 1.5% MB agarose pad and incubated in a temperature controlled chamber (Pecon-Zeiss) at 37°C. Samples were photographed using Axio Observer Z1 (Zeiss) or Eclipse Ti microscope (Nikon, Japan), equipped with CoolSnap HQII camera (Photometrics, Roper Scientific, USA). System control and image processing were performed using MetaMorph 7.4 software (Molecular Devices) or NIS Elements AR 4.3 (Nikon, Japan).

### Live imaging of developing plaques

A custom designed construct (Mamou et al., 2016) was used to monitor bacterial plaques by confocal microscopy. Accordingly, a 40 mm metal ring was filled with 1.5% MB agarose and assembled, and the bacterial cells were infected at low MOI and spotted over the agarose pad. Plaque growth construct was covered with a 35 mm cultFoil membrane (Pecon) to reduce agar dehydration, and incubated in Lab-Tek S1 heating insert (Pecon) placed inside an incubator XL-LSM 710 S1 (Pecon). Initial plaques could be observed under the microscope at t=4 hr. Developing plaques were visualized and photographed by CLSM LSM700 (Zeiss). Cells expressing GFP or YFP were irradiated using 488 nm laser beam, while mCherry expressing cells were irradiated using 555 nm laser beam. For each experiment, both transmitted and reflected light were collected from 6 consecutive Z positions by 10 µm steps. System control and image processing were carried using Zen software version 5.5 (Zeiss).

### Statistical analysis

Unless stated otherwise, bar charts and graphs display a mean ± SD from at least 3 repeats. Quantifications of CPU, PFU, Plaque diameter, infected cells were done manually. MS Excel was used for all statistical analysis, data processing, and presentation.

---

**Table S1.**
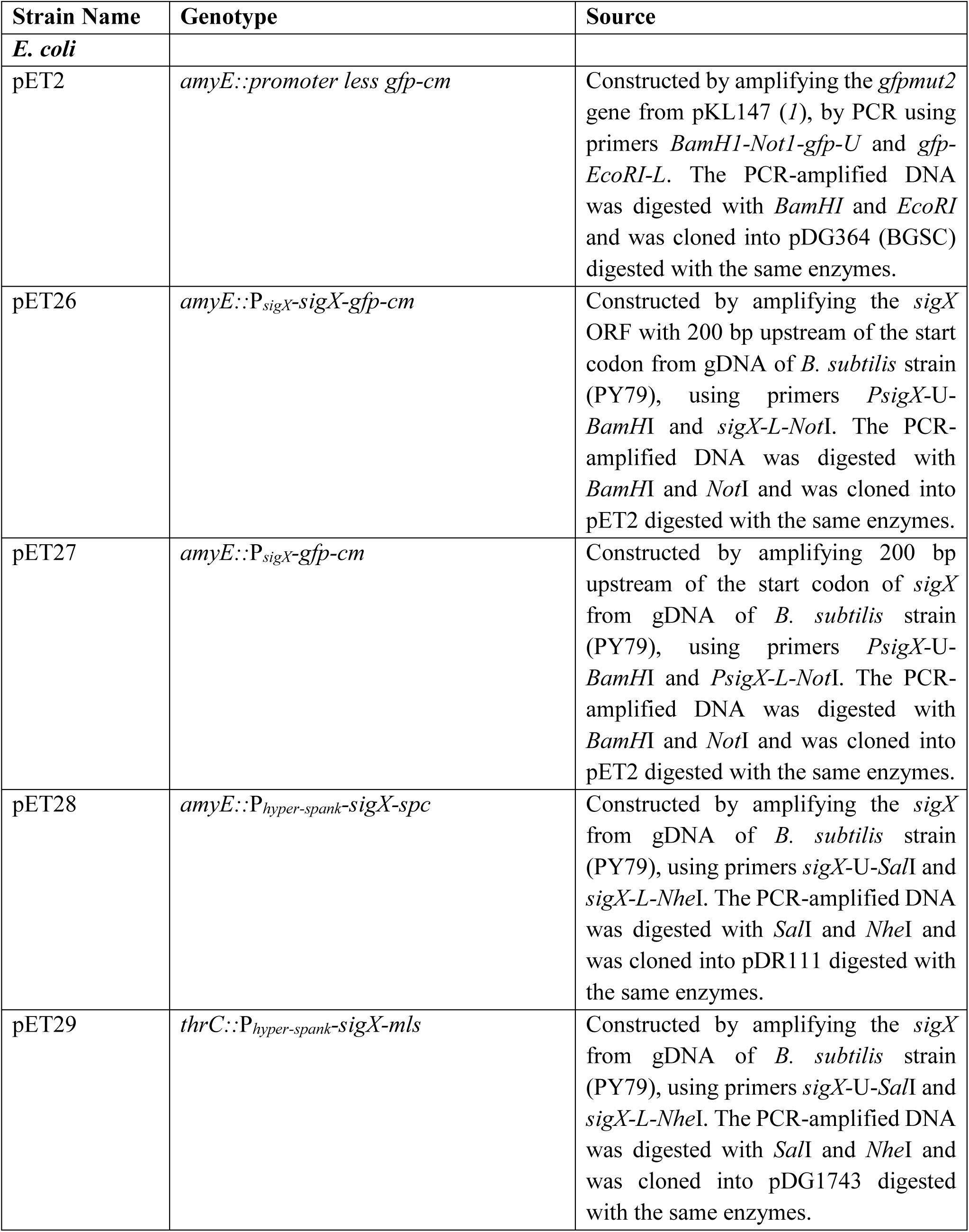
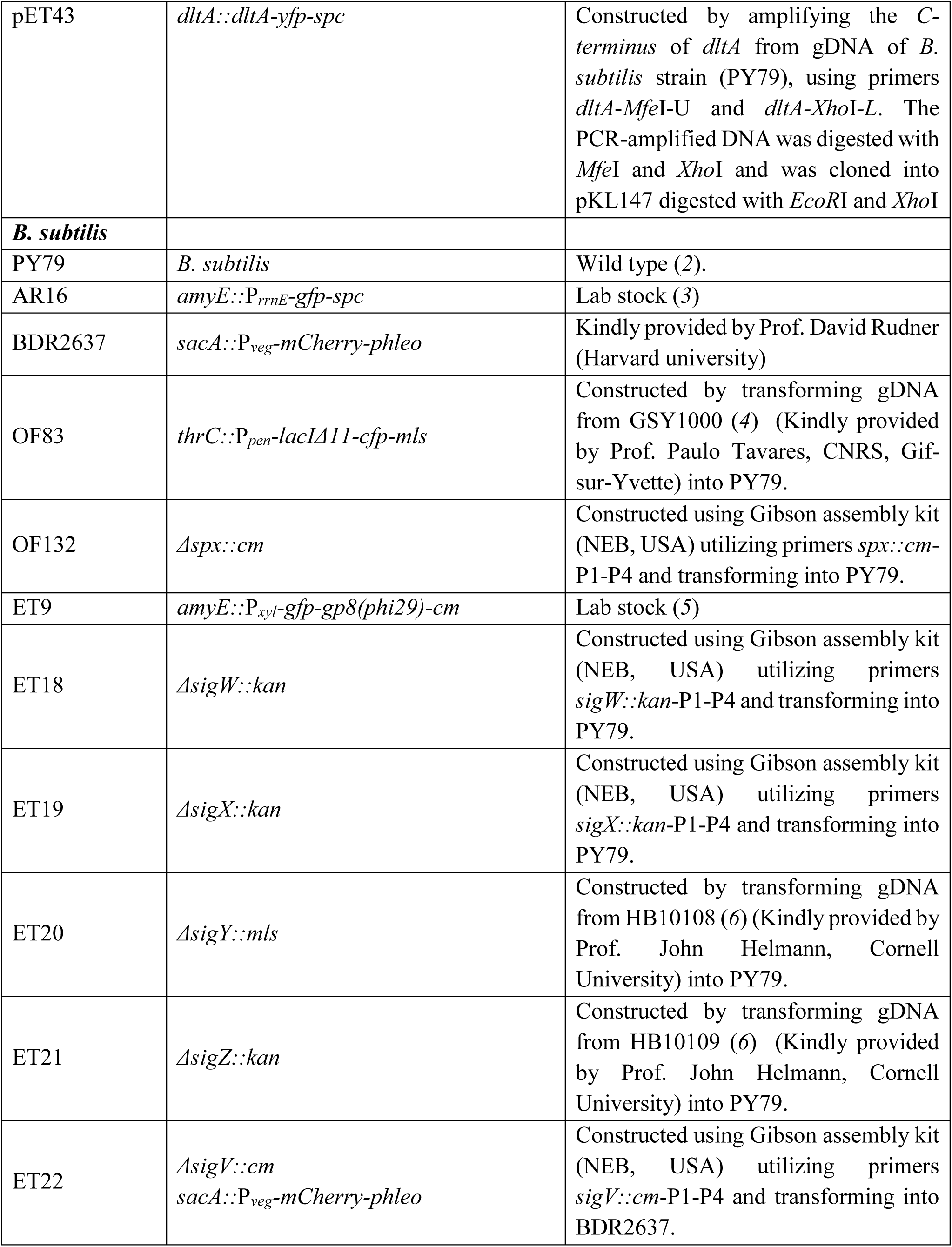
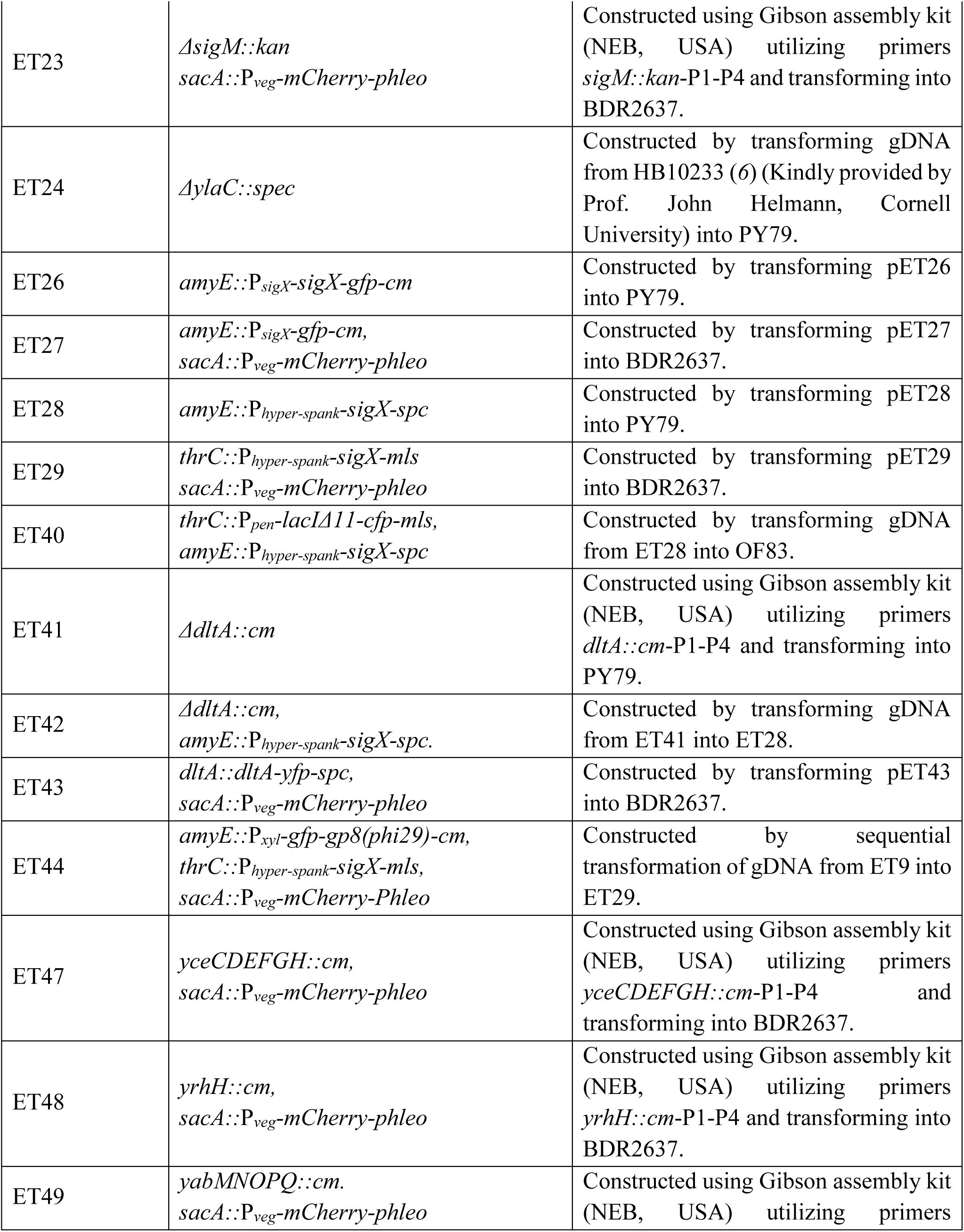
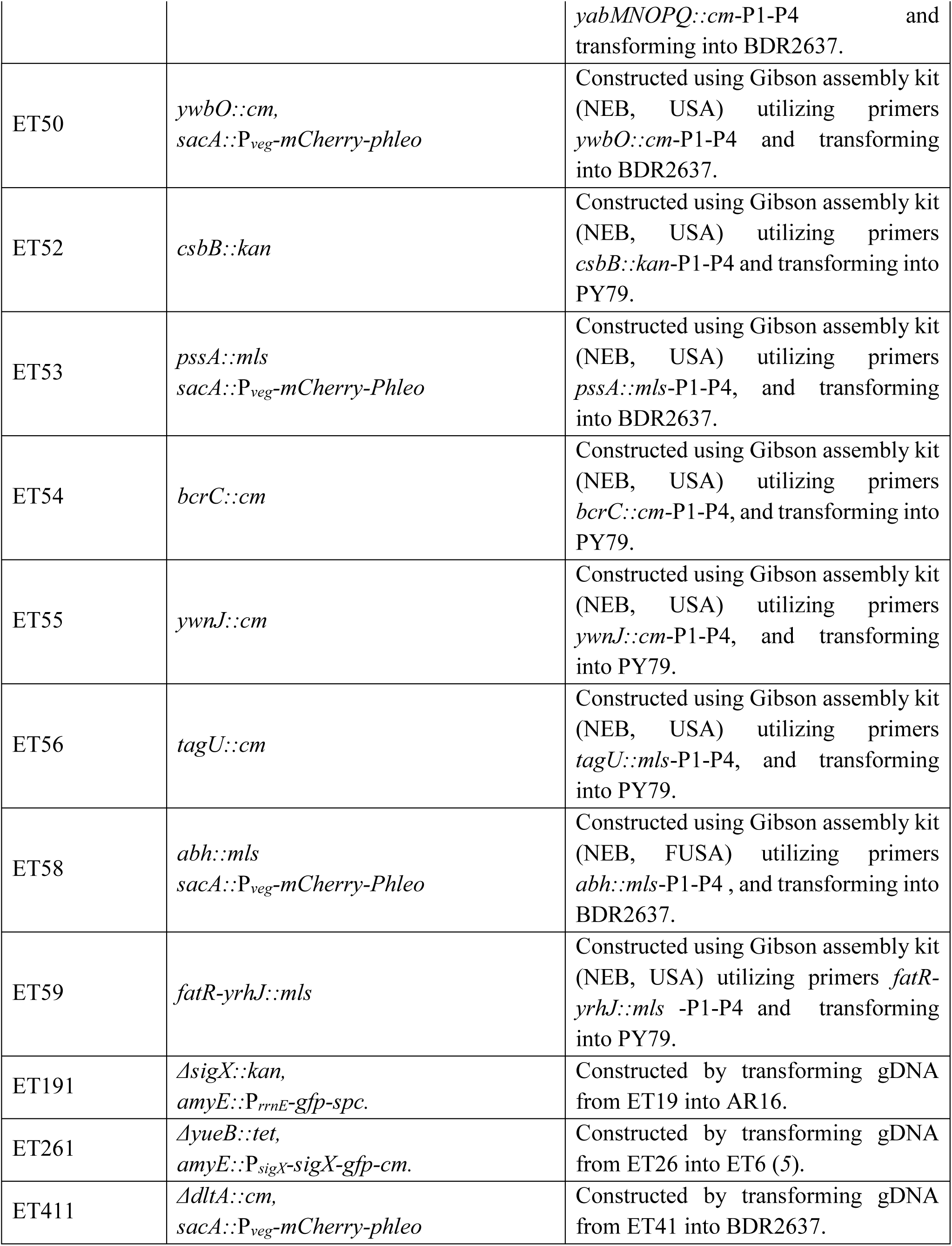
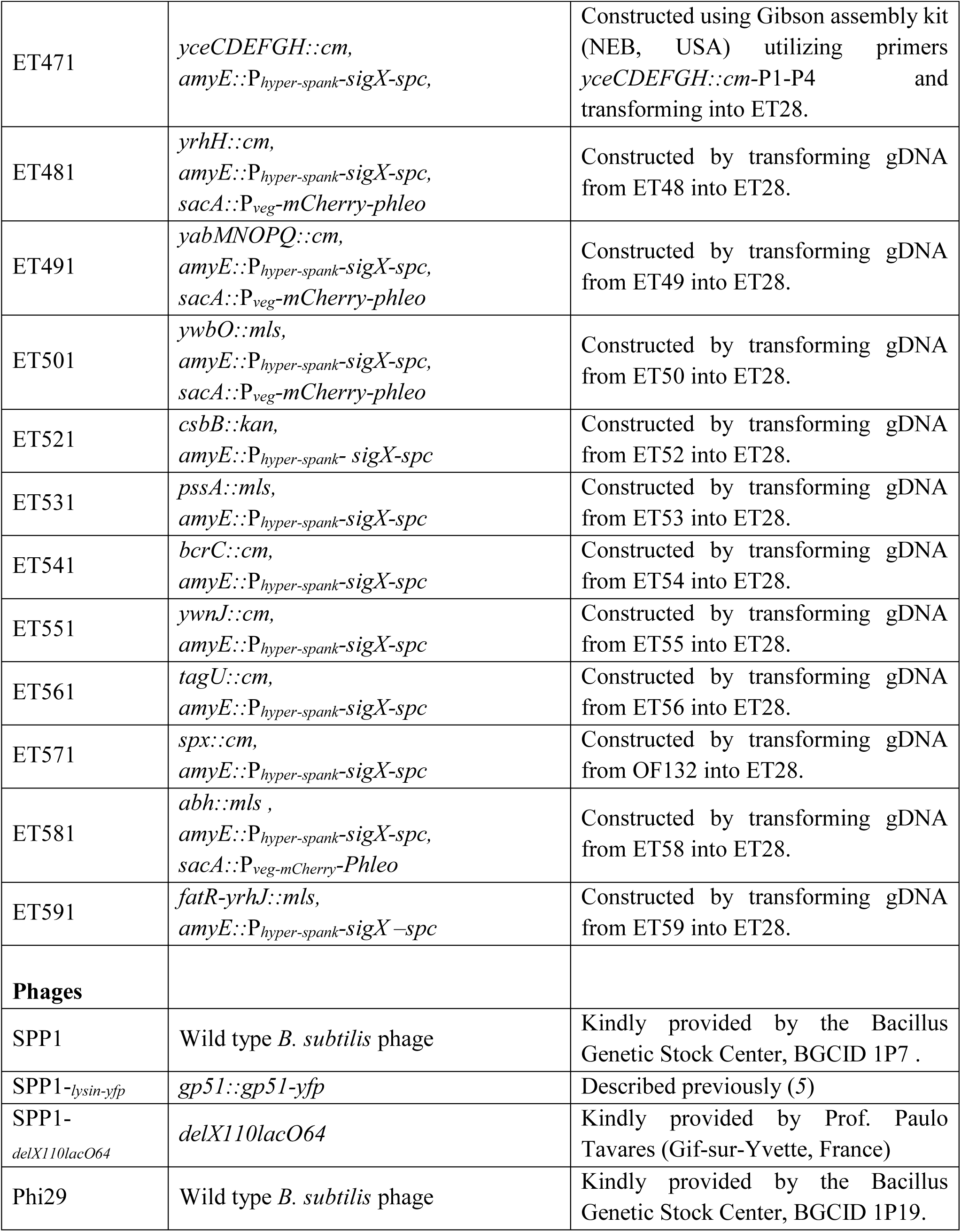
List of bacterial strains and phages used in this study.

**Table S2.**
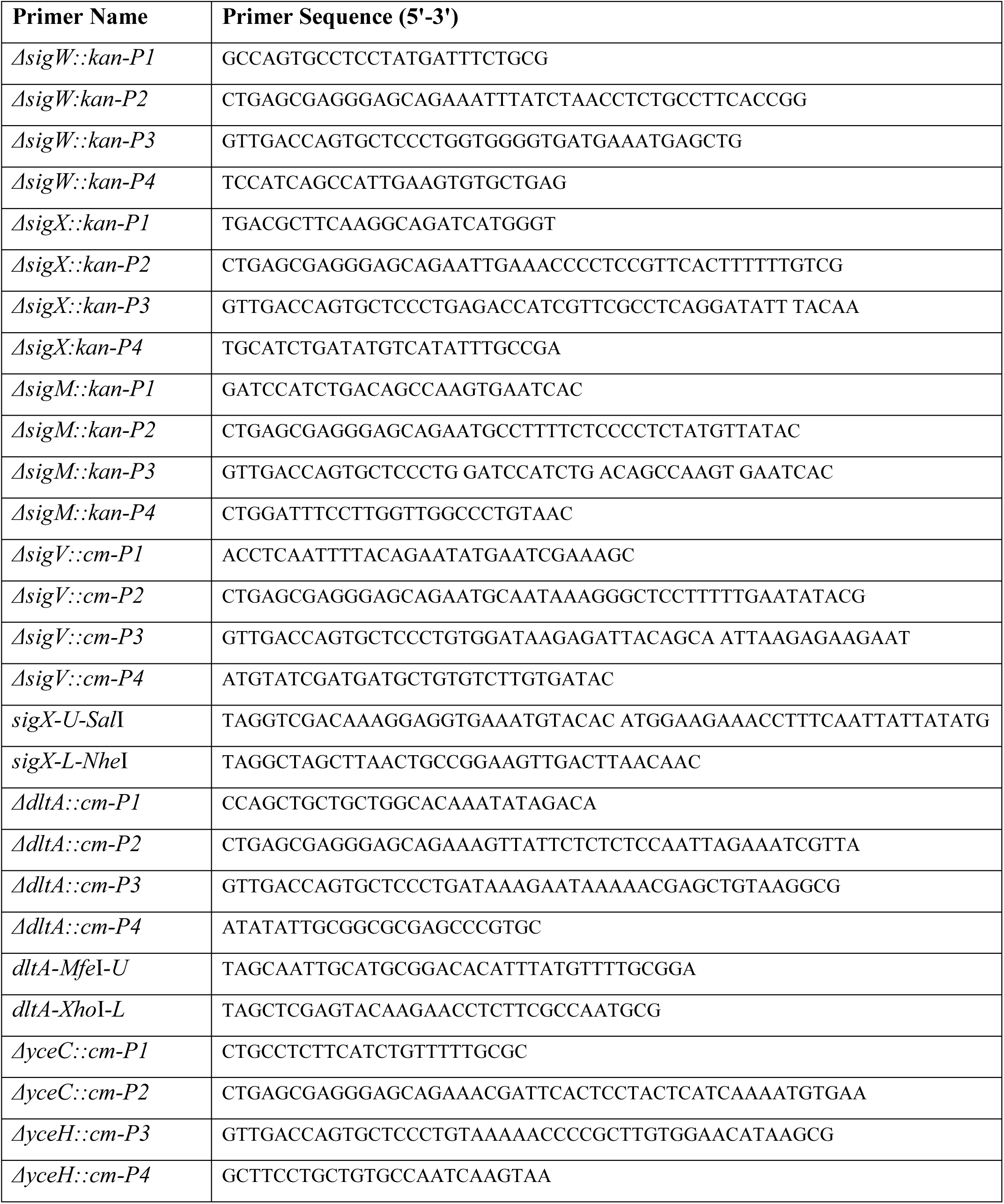
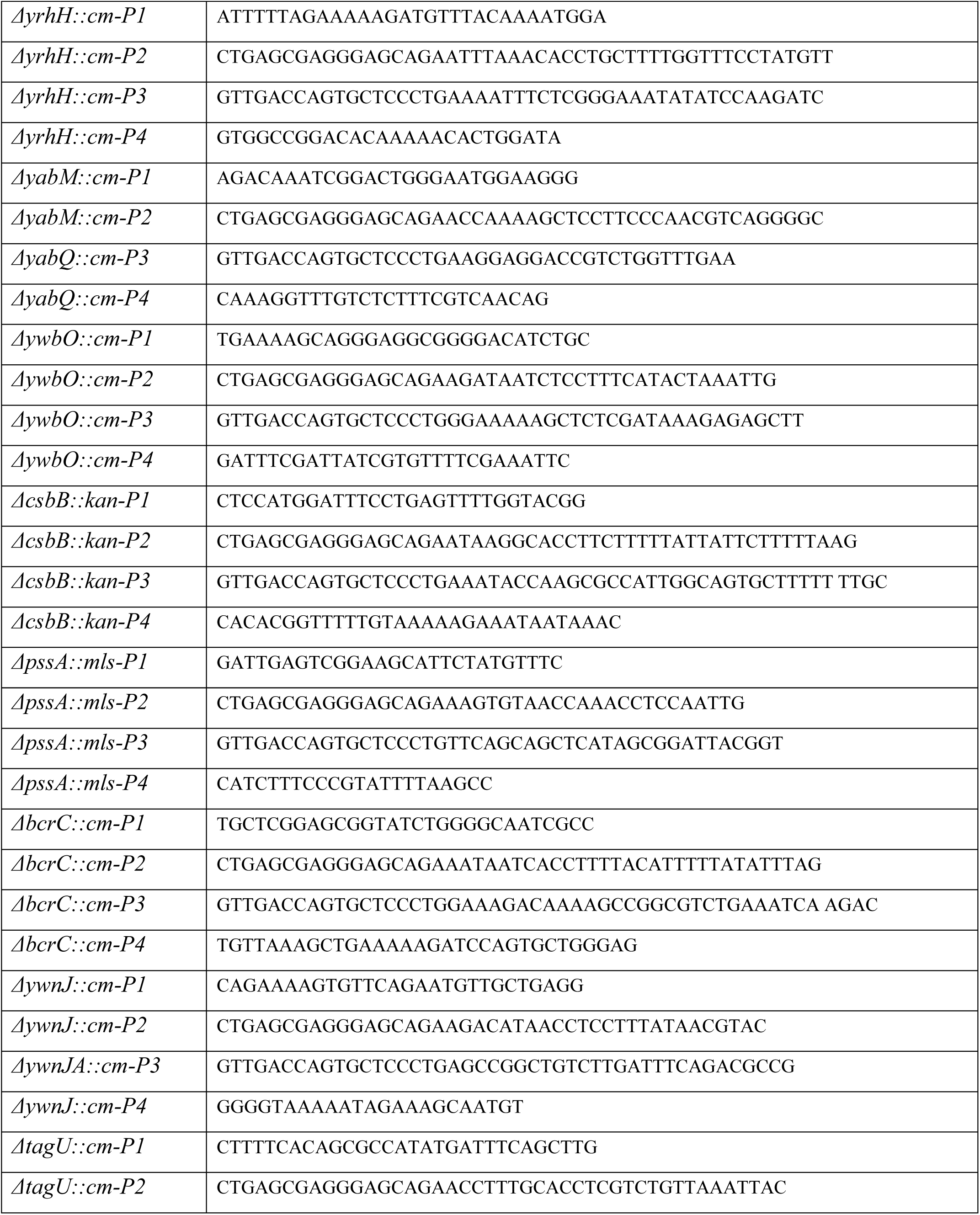
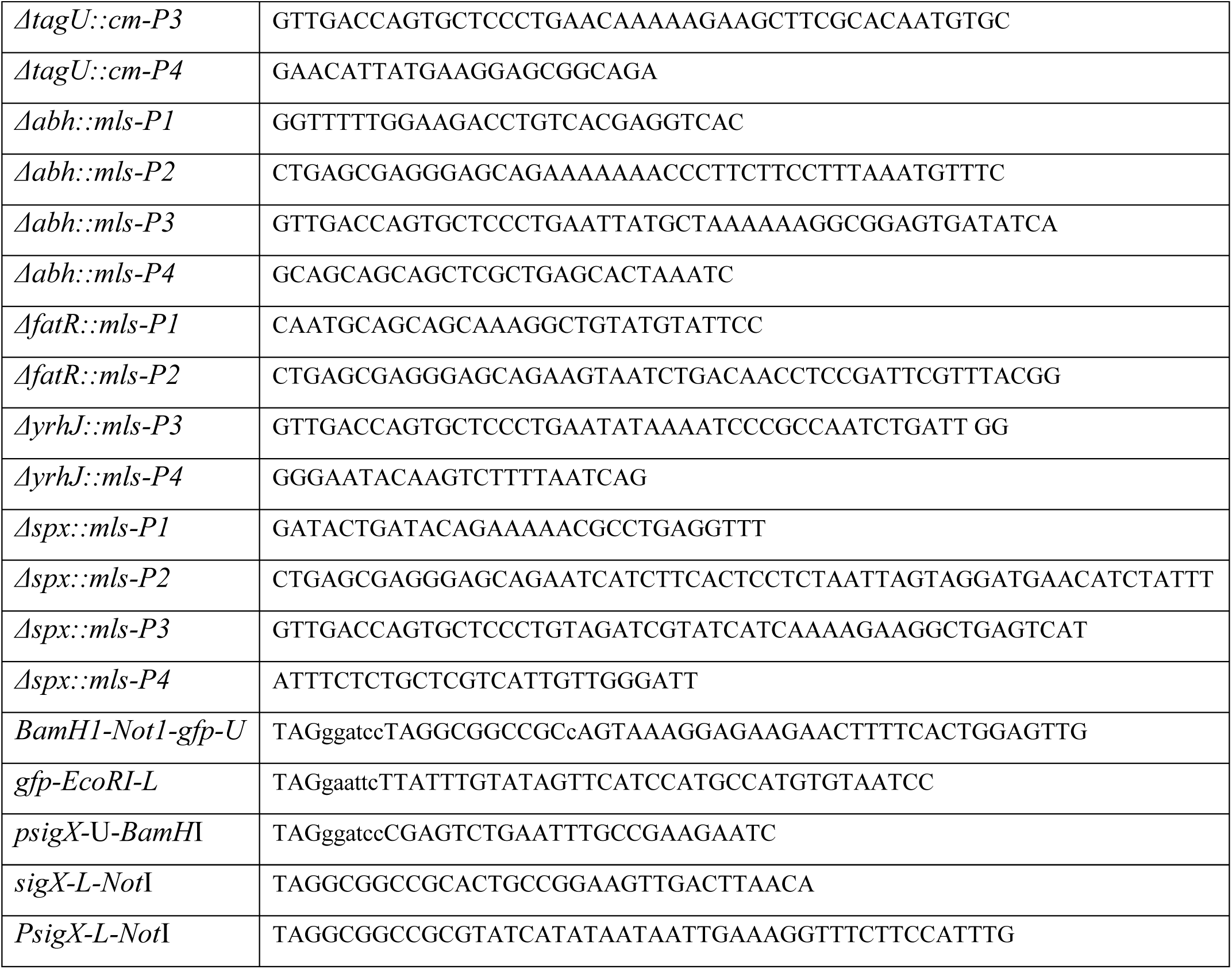
List of primers used in this study.

## References

1. Abedon, S.T., and Yin, J. (2009). Bacteriophage plaques: theory and analysis. Methods Mol Biol 501, 161–174.

2. Alonso, J.C., Luder, G., Stiege, A.C., Chai, S., Weise, F., and Trautner, T.A. (1997). The complete nucleotide sequence and functional organization of *Bacillus subtilis* bacteriophage SPP1. Gene 204, 201–212.

3. Baptista, C., Santos, M.A., and Sao-Jose, C. (2008). Phage SPP1 reversible adsorption to *Bacillus subtilis* cell wall teichoic acids accelerates virus recognition of membrane receptor YueB. J Bacteriol 190, 4989–4996.

4. Brown, S., Santa Maria, J.P., Jr., and Walker, S. (2013). Wall teichoic acids of gram-positive bacteria. Annu Rev Microbiol 67, 313–336.

5. Cao, M., and Helmann, J.D. (2004). The *Bacillus subtilis* extracytoplasmic-function sigmaX factor regulates modification of the cell envelope and resistance to cationic antimicrobial peptides. J Bacteriol 186, 1136–1146.

6. Chanishvili, N. (2012). Phage therapy-history from Twort and d’Herelle through Soviet experience to current approaches. Adv Virus Res 83, 3–40.

7. Cohen, D., Melamed, S., Millman, A., Shulman, G., Oppenheimer-Shaanan, Y., Kacen, A., Doron, S., Amitai, G., and Sorek, R. (2019). Cyclic GMP-AMP signalling protects bacteria against viral infection. Nature 574, 691–695.

8. d’Hérelles, F. (1917). Sur un microbe invisible antagoniste des bacilles dysentériques. Comptes Rendus de l’Académie des Sciences de Paris 165, 373–375.

9. Doron, S., Melamed, S., Ofir, G., Leavitt, A., Lopatina, A., Keren, M., Amitai, G., and Sorek, R. (2018). Systematic discovery of antiphage defense systems in the microbial pangenome. Science 359.

10. Ellis, E.L., and Delbruck, M. (1939). The growth of bacteriophage. J Gen Physiol 22, 365–384.

11. Fernandes, S., Labarde, A., Baptista, C., Jakutyte, L., Tavares, P., and Sao-Jose, C. (2016). A non-invasive method for studying viral DNA delivery to bacteria reveals key requirements for phage SPP1 DNA entry in *Bacillus subtilis* cells. Virology 495, 79–91.

12. Gallucci, S., and Matzinger, P. (2001). Danger signals: SOS to the immune system. Curr Opin Immunol 13, 114–119.

13. Gao, L., Altae-Tran, H., Bohning, F., Makarova, K.S., Segel, M., Schmid-Burgk, J.L., Koob, J., Wolf, Y.I., Koonin, E.V., and Zhang, F. (2020). Diverse enzymatic activities mediate antiviral immunity in prokaryotes. Science 369, 1077–1084.

14. Gottesman, S. (2019). Trouble is coming: Signaling pathways that regulate general stress responses in bacteria. J Biol Chem 294, 11685–11700.

15. Habusha, M., Tzipilevich, E., Fiyaksel, O., and Ben-Yehuda, S. (2019). A mutant bacteriophage evolved to infect resistant bacteria gained a broader host range. Mol Microbiol 111, 1463–1475.

16. Hampton, H.G., Watson, B.N.J., and Fineran, P.C. (2020). The arms race between bacteria and their phage foes. Nature 577, 327–336.

17. Heil, M., and Land, W.G. (2014). Danger signals - damaged-self recognition across the tree of life. Front Plant Sci 5, 578.

18. Helmann, J.D. (2002). The extracytoplasmic function (ECF) sigma factors. Adv Microb Physiol 46, 47–110.

19. Helmann, J.D. (2016). *Bacillus subtilis* extracytoplasmic function (ECF) sigma factors and defense of the cell envelope. Curr Opin Microbiol 30, 122–132.

20. Ho, T.D., and Ellermeier, C.D. (2012). Extra cytoplasmic function sigma factor activation. Curr Opin Microbiol 15, 182–188.

21. Huang, X., Decatur, A., Sorokin, A., and Helmann, J.D. (1997). The *Bacillus subtilis* sigma(X) protein is an extracytoplasmic function sigma factor contributing to survival at high temperature. J Bacteriol 179, 2915–2921.

22. Huang, X., and Helmann, J.D. (1998). Identification of target promoters for the *Bacillus subtilis* sigma X factor using a consensus-directed search. J Mol Biol 279, 165–173.

23. Ingmer, H., Gerlach, D., and Wolz, C. (2019). Temperate phages of *Staphylococcus aureus*. Microbiol Spectr 7.

24. Isaacs, A., and Lindenmann, J. (1957). Virus interference. I. The interferon. Proc R Soc Lond B Biol Sci 147, 258–267.

25. Jakutyte, L., Lurz, R., Baptista, C., Carballido-Lopez, R., Sao-Jose, C., Tavares, P., and Daugelavicius, R. (2012). First steps of bacteriophage SPP1 entry into *Bacillus subtilis*. Virology 422, 425–434.

26. Kingston, A.W., Liao, X., and Helmann, J.D. (2013). Contributions of the sigma(W), sigma(M) and sigma(X) regulons to the lantibiotic resistome of *Bacillus subtilis*. Mol Microbiol 90, 502–518.

27. Kovacs, M., Halfmann, A., Fedtke, I., Heintz, M., Peschel, A., Vollmer, W., Hakenbeck, R., and Bruckner, R. (2006). A functional *dlt* operon, encoding proteins required for incorporation of D-alanine in teichoic acids in gram-positive bacteria, confers resistance to cationic antimicrobial peptides in *Streptococcus pneumoniae*. J Bacteriol 188, 5797–5805.

28. Labrie, S.J., Samson, J.E., and Moineau, S. (2010). Bacteriophage resistance mechanisms. Nat Rev Microbiol 8, 317–327.

29. Levin-Reisman, I., Gefen, O., Fridman, O., Ronin, I., Shwa, D., Sheftel, H., and Balaban, N.Q. (2010). Automated imaging with ScanLag reveals previously undetectable bacterial growth phenotypes. Nature methods 7, 737–739.

30. Lindberg, A.A. (1973). Bacteriophage receptors. Annu Rev Microbiol 27, 205–241.

31. Ma, D., Wang, Z., Merrikh, C.N., Lang, K.S., Lu, P., Li, X., Merrikh, H., Rao, Z., and Xu, W. (2018). Crystal structure of a membrane-bound O-acyltransferase. Nature 562, 286–290.

32. Makarova, K.S., Wolf, Y.I., Snir, S., and Koonin, E.V. (2011). Defense islands in bacterial and archaeal genomes and prediction of novel defense systems. J Bacteriol 193, 6039–6056.

33. McNab, F., Mayer-Barber, K., Sher, A., Wack, A., and O’Garra, A. (2015). Type I interferons in infectious disease. Nat Rev Immunol 15, 87–103.

34. Morehouse, B.R., Govande, A.A., Millman, A., Keszei, A.F.A., Lowey, B., Ofir, G., Shao, S., Sorek, R., and Kranzusch, P.J. (2020). STING cyclic dinucleotide sensing originated in bacteria. Nature 586, 429–433.

35. Patterson, A.G., Yevstigneyeva, M.S., and Fineran, P.C. (2017). Regulation of CRISPR-Cas adaptive immune systems. Curr Opin Microbiol 37, 1–7.

36. Percy, M.G., and Grundling, A. (2014). Lipoteichoic acid synthesis and function in gram-positive bacteria. Annu Rev Microbiol 68, 81–100.

37. Perego, M., Glaser, P., Minutello, A., Strauch, M.A., Leopold, K., and Fischer, W. (1995). Incorporation of D-alanine into lipoteichoic acid and wall teichoic acid in *Bacillus subtilis*. Identification of genes and regulation. J Biol Chem 270, 15598–15606.

38. Saar-Dover, R., Bitler, A., Nezer, R., Shmuel-Galia, L., Firon, A., Shimoni, E., Trieu-Cuot, P., and Shai, Y. (2012). D-alanylation of lipoteichoic acids confers resistance to cationic peptides in group B *streptococcus* by increasing the cell wall density. PLoS Pathog 8, e1002891.

39. Salas, M. (2012). My life with bacteriophage phi29. J Biol Chem 287, 44568–44579.

40. Salmond, G.P., and Fineran, P.C. (2015). A century of the phage: past, present and future. Nat Rev Microbiol 13, 777–786.

41. Sao-Jose, C., Baptista, C., and Santos, M.A. (2004). *Bacillus subtilis* operon encoding a membrane receptor for bacteriophage SPP1. J Bacteriol 186, 8337–8346.

42. Simanski, M., Glaser, R., Koten, B., Meyer-Hoffert, U., Wanner, S., Weidenmaier, C., Peschel, A., and Harder, J. (2013). *Staphylococcus aureus* subverts cutaneous defense by D-alanylation of teichoic acids. Exp Dermatol 22, 294–296.

43. Storz, G. (2016). New perspectives: Insights into oxidative stress from bacterial studies. Arch Biochem Biophys 595, 25–27.

44. Sumrall, E.T., Keller, A.P., Shen, Y., and Loessner, M.J. (2020). Structure and function of *Listeria* teichoic acids and their implications. Mol Microbiol 113, 627–637.

45. Twort, F.W. (1914). An investigation on the nature of ultra-microscopic viruses. Lancet 186, 1241–1243.

46. Tzipilevich, E., Habusha, M., and Ben-Yehuda, S. (2017). Acquisition of phage sensitivity by bacteria through exchange of phage receptors. Cell 168, 186–199 e112.

47. Young, F.E. (1967). Requirement of glucosylated teichoic acid for adsorption of phage in *Bacillus subtilis 168*. Proc Natl Acad Sci U S A 58, 2377–2384.

48. Young, J.C., Dill, B.D., Pan, C., Hettich, R.L., Banfield, J.F., Shah, M., Fremaux, C., Horvath, P., Barrangou, R., and Verberkmoes, N.C. (2012). Phage-induced expression of CRISPR-associated proteins is revealed by shotgun proteomics in *Streptococcus thermophilus*. PLoS One 7, e38077.

## References

Ellis, E.L., and Delbruck, M. (1939). The growth of bacteriophage. J Gen Physiol 22, 365–384.

Levin-Reisman, I., Gefen, O., Fridman, O., Ronin, I., Shwa, D., Sheftel, H., and Balaban, N.Q. (2010). Automated imaging with ScanLag reveals previously undetectable bacterial growth phenotypes. Nature methods 7, 737–739.

Mamou, G., Malli Mohan, G.B., Rouvinski, A., Rosenberg, A., and Ben-Yehuda, S. (2016). Early Developmental Program Shapes Colony Morphology in Bacteria. Cell reports 14, 1850–1857.

Youngman, P., Perkins, J.B., and Losick, R. (1984). A novel method for the rapid cloning in Escherichia coli of *Bacillus subtilis* chromosomal DNA adjacent to Tn917 insertions. Molecular & general genetics : MGG 195, 424–433.

## References

1. K. P. Lemon, A. D. Grossman, Localization of bacterial DNA polymerase: evidence for a factory model of replication. Science 282, 1516–1519 (1998).

2. P. Youngman, J. B. Perkins, R. Losick, A novel method for the rapid cloning in Escherichia coli of *Bacillus subtilis* chromosomal DNA adjacent to Tn917 insertions. Mol Gen Genet 195, 424–433 (1984).

3. A. Rosenberg, L. Sinai, Y. Smith, S. Ben-Yehuda, Dynamic expression of the translational machinery during *Bacillus subtilis* life cycle at a single cell level. PLoS One 7, e41921 (2012).

4. L. Jakutyte et al., Bacteriophage infection in rod-shaped gram-positive bacteria: evidence for a preferential polar route for phage SPP1 entry in *Bacillus subtilis*. J Bacteriol 193, 4893–4903 (2011).

5. E. Tzipilevich, M. Habusha, S. Ben-Yehuda, Acquisition of phage sensitivity by bacteria through exchange of phage receptors. Cell 168, 186–199 e112 (2017).

6. T. Mascher, A. B. Hachmann, J. D. Helmann, Regulatory overlap and functional redundancy among *Bacillus subtilis* extracytoplasmic function sigma factors. J Bacteriol 189, 6919–6927 (2007).

